# Single-cell multiomics identifies key nodes and cis-regulatory elements of the networks specifying the eye domains in zebrafish

**DOI:** 10.64898/2026.01.31.703039

**Authors:** Javier Macho Rendón, Rocío Polvillo, Álvaro Gónzalez-Cid, Jorge Corbacho, Silvia Naranjo, Sofia Manzo, Ana Sousa-Ortega, Ana Fernández-Miñán, Juan Tena, Juan Ramón Martínez-Morales

**Affiliations:** Centro Andaluz de Biología del Desarrollo (CSIC/UPO/JA). 41013 Sevilla, Spain

**Keywords:** Single-cell multiome, gene regulatory networks, optic cup, zebrafish, CRISPR-Cas9, gene network connectivity, network inference, MatchaiRen, *foxp1*

## Abstract

Congenital eye malformations result from disruptions in the early developmental programs that specify and regionalize the eye primordium. Human genetics and developmental studies in animal models have identified critical regulatory nodes within the gene regulatory networks (GRNs) patterning the distinct eye domains. However, the fundamental cis-regulatory connectivity of these networks, and their dynamic behaviour, remains poorly understood. To address these gaps, we performed a comprehensive analysis of transcriptome dynamics and chromatin accessibility at single-cell resolution in the developing zebrafish forebrain. Our single-cell multiomic approach enabled us to characterize the regulatory landscape of the main eye derivatives: neural retina, optic stalk, pigmented epithelium and lens; alongside adjacent diencephalic and telencephalic regions. We identified and functionally validated tissue-specific cis-regulatory modules, and predicted highly connected nodes within each domain. Notably, we uncovered the role of *foxp1b* as a conserved central node within the neural retina network, highlighting the predictive power of our analysis in reconstructing the architecture of the GRNs that specify the vertebrate eye.

**Teaser:** Identification of central nodes and cis-regulatory networks involved in eye patterning by single-cell multiomics in zebrafish.

## Introduction

The vertebrate brain blueprint emerged more than 500 million years ago in the common ancestor of our phylum. Despite the broad variety of morphological and physiological adaptations in vertebrate brains, this shared origin is reflected in a preserved basic scaffold. This becomes especially evident during early embryogenesis, particularly at the phylotypic period, when the developing brain converges into the prosomeric organization recognizable across all species (*1*). Within this general plan, the eye primordium is specified during neurulation in the anterior neuroectoderm, as distinct forebrain domains (namely the telencephalic, retinal, and diencephalic territories) are patterned along the anteroposterior and dorsoventral axes under the influence of signalling molecules secreted from organizing centres (*2*). Although substantial molecular evidence indicates that the main nodes of the GRNs specifying the vertebrate forebrain are conserved even at the chordate level (*3*), our understanding of their precise architecture remains fragmentary, especially at the cis-regulatory level, and species-specific differences have been reported (*4*). In this broader context, the GRNs involved in eye specification are particularly well conserved within vertebrates, consistent with the fact that both the ontogeny and final architecture of the eye are well preserved, even in primitive jawless vertebrates (*5*).

Over the past few decades, the central nodes comprising the eye networks have been identified primarily through genetic analyses in vertebrate models, including mice, chicken, *Xenopus*, zebrafish and medaka (*6*). Many of these key nodes have also been identified as causative genes in common congenital eye malformations within the spectrum of anophthalmia, microphthalmia and coloboma syndromes (MACs). These severe eye disorders are a frequent cause of visual impairment in children, with a combined prevalence of up to 3 in 10,000 births (*7, 8*). MAC-causing genes predominantly encode transcriptional regulators such as *SOX2*, *OTX2*, *RAX*, *FOXE3*, *PAX6*, *VSX2*, *MAF*, *SIX3*, *MITF*, *PITX3* or *YAP1*. In addition, the list includes genes encoding for a wide variety of molecules involved in growth factor signalling, retinoic acid metabolism, extracellular matrix components, intracellular trafficking, and other pathways (e.g. *SHH*, *BMP4*, *BMP7*, *GDF3*, *STRA6*, *ALDH1A3*, *COL4A1*, *FRAS*, *RAB18*, or *MAB21L2*) (*9*). As this gene list includes tissue-identity regulators for each of the main territories comprising the eye primordium (i.e. neural retina, RPE, optic stalk, and lens), it is likely that developmental failure in the specification of any of these components may result in a MAC phenotype. Although this group of diseases has a strong genetic component, and causative mutations have been identified for more than 90 genes, the genetic basis of MAC remains unknown in a substantial proportion of patients: estimated at 70-80% (*9*). Even among the more severe cases, such as bilateral anophthalmia, approximately 40% of the patients remain genetically undiagnosed (*10*). Unidentified causative genes, rare variants, complex genetic interactions, and environmental factors may contribute to this hidden heritability. Additionally, there is increasing recognition of the role that non-coding variants may play as pathogenic mutations, especially when these regions show evolutionary conservation across vertebrates (*11, 12*).

Hidden heritability in patients with MACs raises the question of how well we understand the architecture of the gene networks that govern eye specification. The identification rate of central nodes of these networks, through gene-centric studies in vertebrate models or human MAC patients, has declined in recent years. While this may suggest that most key regulators (primarily transcription factors) have already been uncovered, several aspects may hinder the identification of the remaining ones through traditional genetic approaches. Retinal specification networks have been proved to be particularly robust, as parallel transcription factors converge on similar DNA binding motifs (*13*). As a result, the development of the organ can often withstand mutations in individual components, even when they play a central role in the network, such is the case for *vsx* genes mutation in zebrafish (*14*). In teleost models in particular, gene compensation and compensatory growth of the organ may also result in no overt eye phenotype (*15, 16*). Conversely, mutations in pleiotropic genes may lead to early embryonic lethality, thereby obscuring their specific contributions to eye development (*17, 18*).

To understand the architecture of eye networks and predict their behaviour in both normal and pathological contexts, it is essential to characterize not only their nodes but also their connecting edges. These edges represent gene interactions mediated by cis-regulatory elements (CREs), including promoters and enhancers, that control cell-type-specific gene expression. While many of the retinal CREs have been characterized by transgenesis assays, particularly in mice and zebrafish (*19, 20*), only a limited number have been functionally validated through targeted deletion (*21–25*). Beyond genetic approaches, the emergence of next-generation sequencing technologies and single-cell methods has enabled a systems-level perspective on retinal network connectivity (*26*). Single-cell transcriptomics (scRNAseq) has overcome the limitations of bulk RNA profiling for the characterization of heterogeneous cell populations. These studies have facilitated the mapping of conserved and species-specific regulatory modules and differentiation trajectories for all retinal cell types across vertebrates models, including mice, zebrafish, chicken, and human embryos, as well as in human retinal organoids (*27–33*). Moreover, the integration of single-nuclei ATAC-seq with single-nucleus transcriptomic data now enables the use of computational methods to infer GRNs at an unprecedent resolution (*34*). In the context of retinal differentiation, single-cell multimodal profiling has recently been applied to characterize chromatin landscapes and predict the role of relevant enhancers (*35, 36*); as well as to model the GRNs governing neurogenic progenitors specification and neuronal cell types differentiation (*37, 38*).

In this study, we employed a single-cell multiome approach that complements prior work focused on retinal neuronal subtype specification by shifting the focus to earlier stages of zebrafish eye development. We characterized the regulatory landscape of eye domains as their identities are established, including the neural retina, optic stalk, retinal pigmented epithelium, and lens placode, and extended our analysis to adjacent forebrain territories such as the telencephalon and diencephalon. To this end we retrieved transcriptomic information from 20,763 individual cells at 15, 18, and 23 hours post-fertilization (hpf), and integrated transcriptomics and chromatin accessibility from 14,582 nuclei at 18 hpf. Our analysis enabled the reconstruction of tissue-specific transcriptomic dynamics, the identification and validation of highly-specific cis-regulatory modules, and the discovery of central regulatory nodes within each GRN. As proof of principle of the predictive power of our approach, we identified *foxp1b* as a conserved hub within the neural retina GRN and validated its function through CRISPR-Cas9 mediated deletion. This work provides new insights into the organizational logic of the regulatory networks of the eye and may serve as a valuable reference for investigating the aetiology of inherited eye malformations.

## Results

### Analysis of zebrafish eye and forebrain specification networks by sc-RNA-seq

To gain insight into the developmental programs specifying the different domains of the eye primordium and adjacent forebrain tissues, we conducted scRNA-seq experiments at three developmental stages in zebrafish —15, 18 and 23 hpf— spanning the transition from optic vesicle to optic cup (Fig 1A). During this period, the identity of the distinct eye domains is established, and neurogenesis has not yet commenced in the neural retina (*13*). Leveraging the broad expression of the *Tg(rx3:Gal4;UAS:GFP)* transgene in the anterior neural plate (*39*), we isolated cells from the eye primordium and adjacent forebrain tissues by FACS, employing a relaxed gating strategy (see Methods). We generated libraries using the 10X Genomics Chromium platform for each sorted sample, profiling a total of 7,447 cells at 15 hpf, 9,620 cells at 18 hpf, and 3,696 cells at 23 hpf, with a median of 59,001 reads per cell and 2,366 genes detected per cell, as determined using *CellRanger* (Supplementary Figure S1). As a reference for cell coverage, the zebrafish optic cup has been estimated to contain approximately 2500 cells at 24hpf (*40*). Detailed quality control metrics, including per-cell read distribution and feature counts, are provided in Supplementary Figure S1.

**Figure 1.**
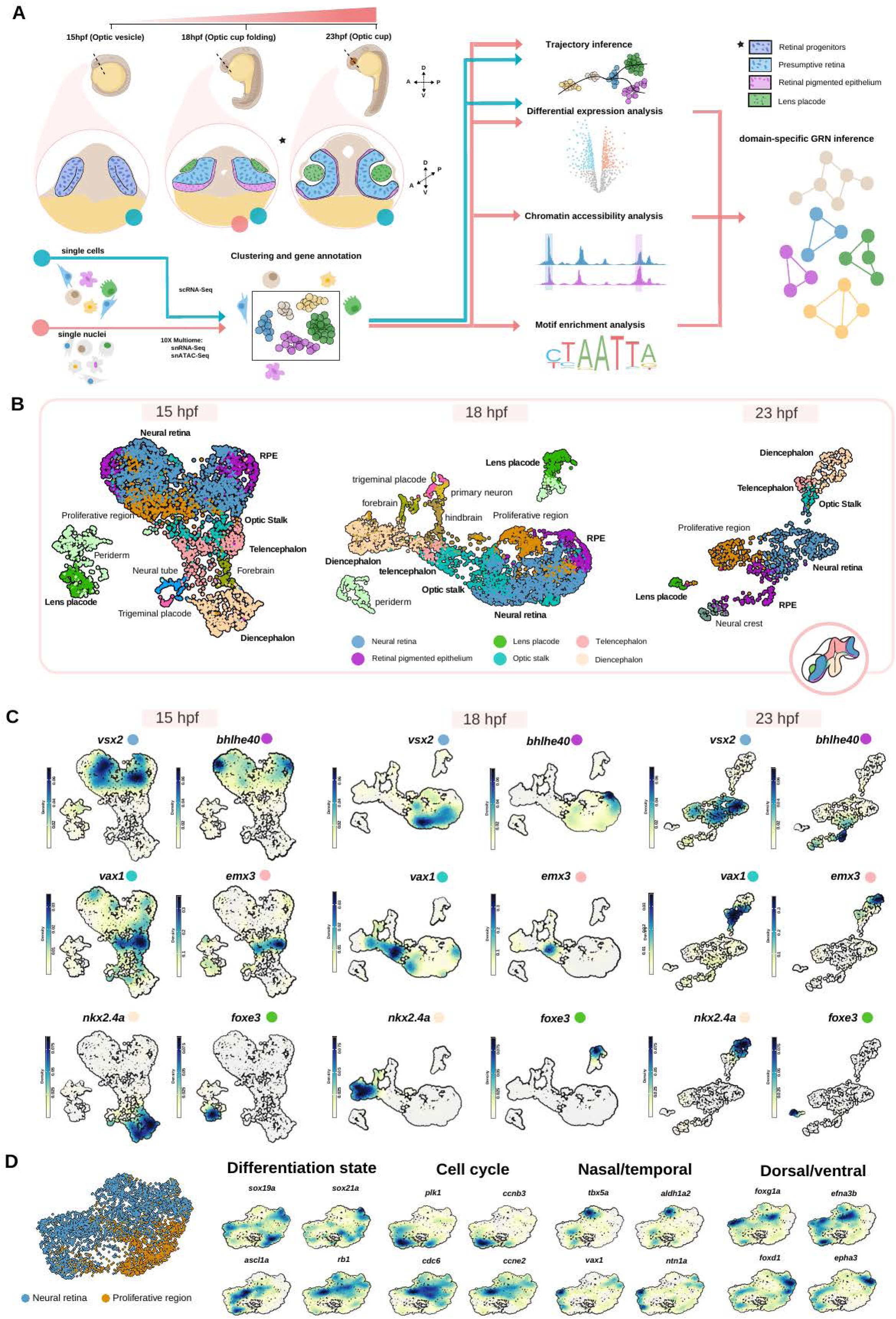
Single-cell transcriptomic and multiome profiling reveal the main domains and transcriptional states of the developing zebrafish eye and adjacent forebrain tissues. **(A)** Overview of the experimental design. Anterior neural plate and eye primordium of zebrafish embryos were collected at three developmental stages spanning the transition from optic vesicle to optic cup (15, 18 and 23 hpf) and profiled using scRNA-seq. In parallel, 18 hpf embryos were profiled using single-nucleus multiome sequencing (snRNA-seq + snATAC-seq). Computational pipelines were applied to each independent dataset, including clustering, cell-type annotation and differential expression analysis. Chromatin accessibility profiling and motif enrichment analysis were also performed for the 18hpf multiome dataset to infer domain-specific gene regulatory networks. **(B)** UMAP embeddings of scRNA-seq datasets from 15, 18 and 23 hpf, coloured according to annotated cell identity. Six major territories—neural retina, RPE, optic stalk, telencephalon, diencephalon and lens placode—were consistently recovered across developmental stages. **(C)** Density plots showing expression of canonical markers for each domain across the three developmental stages: *vsx2* (neural retina), *bhlhe40* (RPE), *vax1* (optic stalk), *emx3* (telencephalon), *nkx2.4a* (diencephalon) and *foxe3* (lens placode). Marker expression co-localizes with the annotated clusters in (B), confirming the robustness of domain identity throughout this developmental window. **(D)** Sub-clustering and integration of neural retina cells across 15, 18 and 23 hpf. The integrated UMAP reveals two transcriptional states within the neural retina: a proliferative precursor population expressing SoxB family genes (*sox19a, sox21a*) and a proneural population expressing differentiation and cell-cycle exit markers (*ascl1a, rb1*). Density plots also highlight markers of cell-cycle phases (*plk1, ccnb3, cdc6, ccne2*) as well as spatial axes within the retina, including markers of dorso–ventral (*tbx5a/aldh1a2* vs. *vax1/ntn1a*) and naso–temporal (*foxg1a/efna3b* vs. *foxd1/epha3*) patterning.

Unsupervised clustering (*Seurat*), combined with curated annotation based on differentially expressed known markers was used to identify distinct cell clusters. Cluster identities were further refined using enriched anatomical ontology terms associated with DEGs, as annotated in ZFIN (See Methods). A schematic overview of the annotation workflow is provided in Supplementary Figure S1. This analysis delineated the main territories comprising the developing eye and adjacent forebrain regions present at each stage, namely: neural retina, RPE, optic stalk, telencephalon, diencephalon and lens domains, which are color-coded in the UMAP representation shown in Figure 1B. Ontology-guided refinement and clustering also subdivided the neural retina domain into two distinct cell states: a proliferative state (orange in Figure 1B) and a proneural state (blue in Figure 1B), both expressing a shared core set of neural retina markers (Figure 1C). A complete list of DEGs for each territory is provided in Supplementary Dataset 1.

All six major domains were identifiable in each dataset (15, 18 and 23hpf) as confirmed by the expression of the corresponding marker genes: *vsx2*, *bhlhe40*, *vax1*, *nkx2.4a*, *emx3* and *foxe3* (Supplementary Dataset 1) (Figure 1C). This indicates that, despite differences in UMAP topology across stages, the overall composition of cell clusters remains stable within datasets in developmental window. To validate our analysis, we examined the expression patterns of the ten most significant transcription factors (ranked by p-value) in each domain. ZFIN annotations showed that 92% of these transcription factors (55 of 60) are already reported as expressed in their corresponding territories in the literature. A complete list of differentially expressed transcription factors is provided in Supplementary Dataset 2.

To further evaluate the resolution of our single-cell analysis, we focused on the transcriptional signature of the neural retina domain to identify potential subclusters within this territory. Integration of annotated retinal populations across stages revealed clear segregation of markers associated with spatial patterning, cell-cycle progression, and differentiation states (Figure 1D). For instance, cells grouped according to their differentiation state into proliferative precursors (orange), characterized by the expression of *Soxb* family transcription factors (*sox19a* and *sox21a*), and committed neural progenitors (blue), expressing proneural and cell-cycle exit markers (*ascl1a* and *rb1*) (Figure 1D, Supplementary Dataset 1). In addition, cells could also be grouped according to cell-cycle markers, with *plk1* and *ccnb3* marking cells in S phase and *cdc6* and *ccne2* labeling mitotic populations. Furthermore, markers of dorso-ventral patterning (*tbx5a*/*aldh1a2* vs. *vax1*/*ntn1a*) and naso-temporal polarity (*foxg1a*/*efna3b* vs. *foxd1*/*epha3*) were clearly segregated within the embedding, consistent with the established organization of the retinal field.

### Developmental trajectories and RNA velocity reveal temporal ordering of retinal differentiation

To characterize the transcriptional dynamics of the developing forebrain populations, we applied RNA velocity using scVelo dynamical modeling across retinal and adjacent forebrain cells at 15, 18, and 23 hpf (Figure 2A). This approach revealed a broadly consistent pattern of transcriptional flow across stages, which was further confirmed by integrating all three datasets (Figure 2B). The results indicated that a core of cells within the neural retina cluster, corresponding to the proliferative state, exhibits near-zero RNA velocity, indicative of stable transcriptomic stage. From this proliferative state core, transcriptional trajectories flow not only toward committed neural retina populations (Figure 2B) but also toward adjacent RPE, optic stalk, and telencephalic territories. By contrast, the diencephalon contains a similarly stable core of cells that appears largely isolated from other populations, suggesting a distinct and relatively independent transcriptomic program. Among non-neural components, lens and periderm cells form discrete groups, with transcriptional flow indicating a progression from periderm toward lens identity, in line with the known ontogeny of the lens primordium. Collectively, these findings support the model that the retinal proliferative state behaves as a transcriptionally naive hub for multiple forebrain territories, with the exception of diencephalic populations.

**Figure 2.**
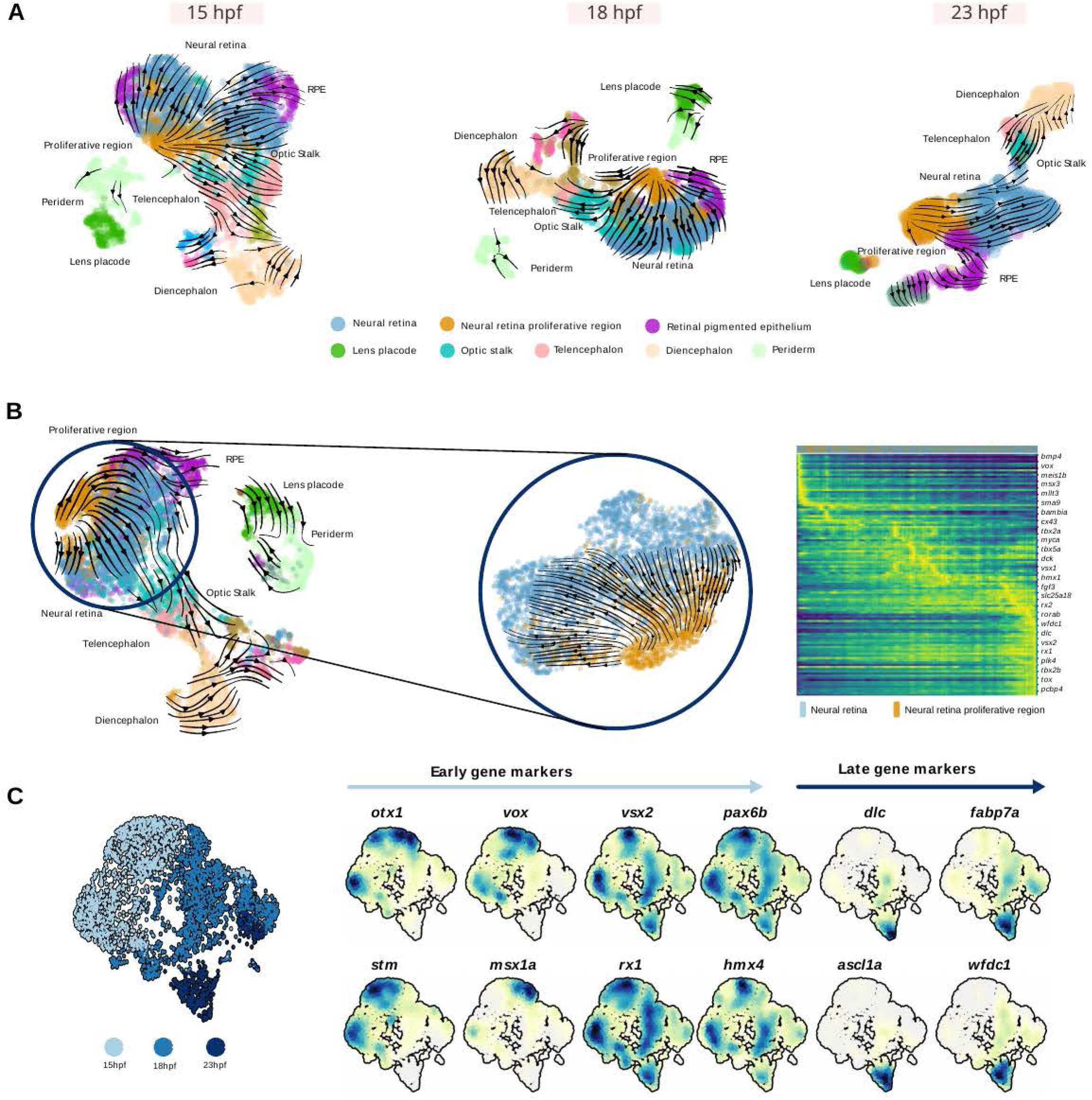
RNA velocity highlights transcriptional dynamics in the developing forebrain populations. **(A)** RNA velocity applied to scRNA-seq datasets at 15, 18 and 23 hpf. UMAP embeddings reproduce the cluster structure shown in Figure 1B, now overlaid with velocity streamlines. **(B)** Integration of the three stages highlights a conserved global flow pattern. Velocity streamlines converge onto the proliferative retinal core and project toward committed neural retina cells as well as neighboring RPE, optic stalk and telencephalic territories. **(C)** Latent time ordering of retinal cells represented as a heatmap of velocity-inferred driver genes. **(D)** Validation of temporal expression in computationally merged retinal cells. The density plots across UMAPs show the progression of representative early markers (*otx1*, *vox*, *stm*, *msx1a*), broadly expressed identity genes (*vsx2*, *pax6b*, *rx1*, *hmx4*) and late/proneural markers (*dlc*, *fabp7a*, *ascl1a*, *wfdc1)*.

To gain deeper insight into the neural retina transcriptional dynamics, we focused on the proliferative and committed states. Within the proliferative population, we observed a heterogeneous pattern of RNA velocity: a subset of cells remained as transcriptionally stable progenitors, whereas others showed clear trajectories toward the committed neural retina state, indicating that a fraction of cycling progenitors is transitioning into a proneural program (Figure 2B). Supporting this interpretation, latent time analysis (an RNA velocity-based measure of transcriptional flow) ordered cells along a continuous proliferative-to-proneural axis and enabled the identification of putative driver genes whose expression dynamics correlate with this transition (Figure 2C). To validate these temporal patterns, we visualized gene expression in datasets merged by developmental stage without forced integration (Figure 2D). This analysis revealed stage-enriched markers, including early regulators (o*tx1, vox, stm, msx1a),* broadly expressed retinal progenitor genes *(vsx2, pax6b, rx1, hmx4)* and late markers associated with proneural commitment and early differentiation *(dlc, fabp7a, ascl1a, wfdc1).* Together, these results reveal a coherent transcriptional progression from proliferative to committed neural retina states, capturing the regulatory transition of early retinal progenitors consistent with the trajectories inferred from RNA velocity.

### Multiomic profiling at 18 hpf integrates transcriptional and chromatin landscapes

Given that the transcriptional identities of the eye and adjacent forebrain populations are highly conserved across the three developmental stages (Figures 1, 2), we next focus on the intermediate stage, 18hpf, to further explore the regulatory logic underlying these cellular identities. To this end, we performed single-nucleus multiome profiling (snRNA-Seq + snATAC-Seq) on 150 dissected embryo heads, capturing both transcriptional profile and chromatin accessibility within individual nuclei. This approach enabled us to link gene expression programs with their regulatory landscapes.

Libraries were generated using the 10X Genomics Chromium platform for each sample, and processed with Cellranger ARC, yielding a total of 14582 nuclei. The median number of reads per nucleus was 33,880 reads for snRNA-Seq (with 1,570 genes per nucleus), and 28,915 reads for snATAC-Seq (with 8049 features per nucleus) (Supplementary Figure S1). A joint dimensionality reduction across modalities approach, combined with our cell annotation method (See Methods), recapitulated the major populations previously observed in scRNA-seq (Figure 3A), as confirmed by the expression of domain-specific beacon markers (*vsx2*, *bhlhe40*, *vax1*, *nkx2.4a*, *emx3*, *foxe3*) (Figure 3B). The only exception was the neural proliferative state cluster, identified in the single-cell datasets (Figure 1B, D), which was not annotated automatically in the multiome profiling. To investigate this, we manually focused on the neural retina cluster and confirmed that markers of spatial patterning, cell-cycle, and differentiation states were also distinguished within the multiome embedding (Supplementary Figure S2).

**Figure 3.**
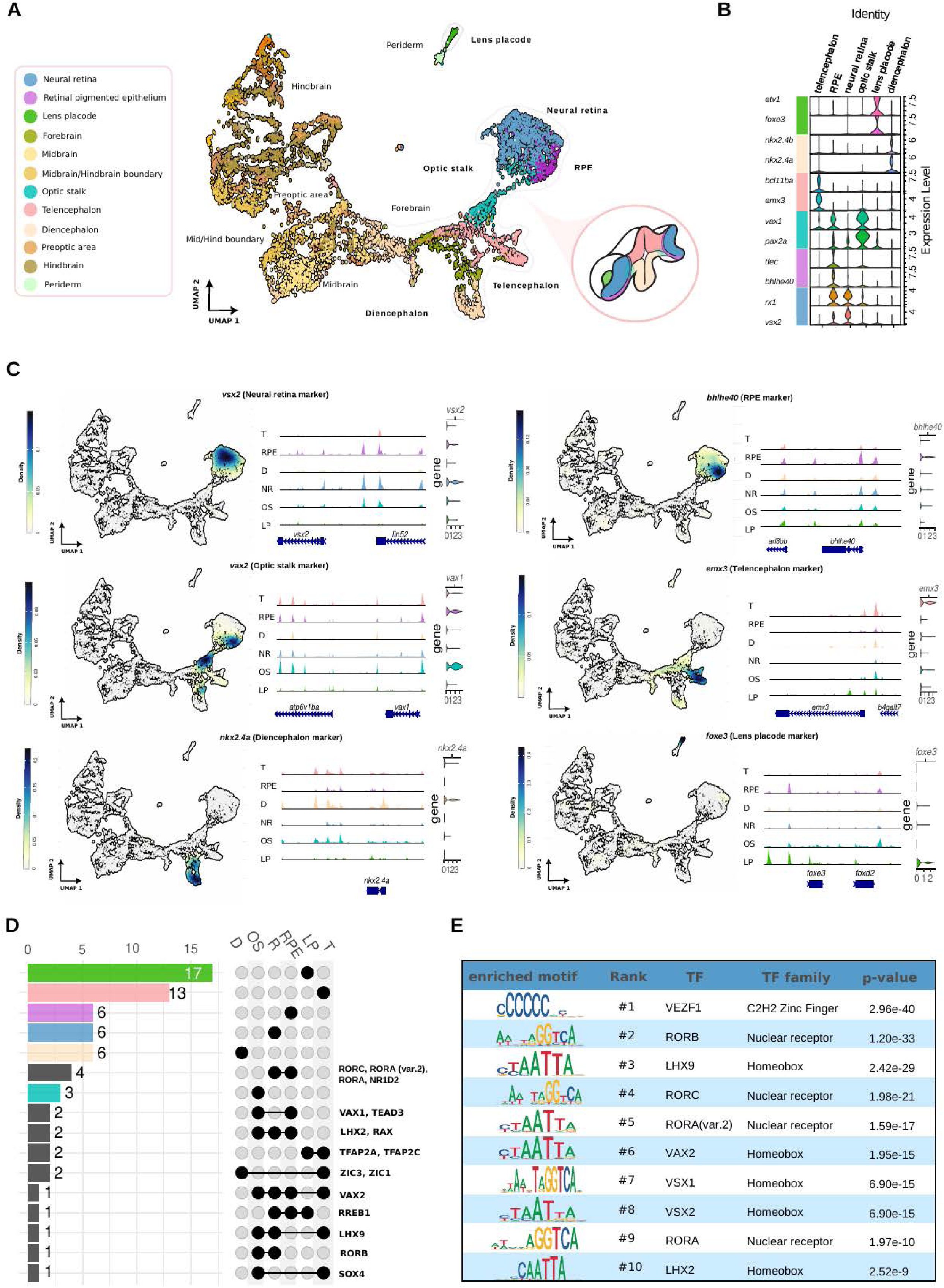
Chromatin accessibility and transcriptional identity across eye and forebrain domains at 18 hpf. **(A)** UMAP of the 18 hpf multiome dataset (snRNA-seq + snATAC-seq), showing major eye and brain territories. The subset corresponding to eye and forebrain populations is highlighted, with domains (neural retina, RPE, lens placode, optic stalk, telencephalon, diencephalon) annotated on a schematic 3D anatomical representation. **(B)** Violin plots showing expression of two canonical markers per domain (*etv1*/*foxe3*: lens placode; *nkx2.4a*/*nkx2.4b*: diencephalon; *bcl11ba*/*emx3*: telencephalon; *vax1*/*pax2a*: optic stalk; *tfec*/*bhlhe40*: RPE; *rx1*/*vsx2*: neural retina). **(C)** Density plots of one domain marker per territory (*vsx2*, *bhlhe40*, *vax1*, *emx3*, *nkx2.4a*, *foxe3*) together with corresponding chromatin accessibility profiles at nearby regulatory regions. Domain-specific snATAC peaks align with marker-gene expression, illustrating coordinated transcriptional and chromatin signatures. **(D)** UpSet plot showing intersections among sets of the most significant enriched motifs across domains. **(E)** Top enriched motifs in the neural retina domain.

Differential expression (DEGs) and differential chromatin accessibility (DOCRs) analyses were performed for each annotated domain, and the resulting lists are provided in Supplementary Dataset 3. We then correlated the expression of beacon markers with cell-type-specific chromatin accessibility at proximal regulatory regions (Figure 3C). These paired RNA-Seq–ATAC-Seq profiles highlight the coordination between transcriptional activity and cis-regulatory accessibility within the same cells. To uncover potential transcriptional regulators, we performed motif enrichment analysis on the DOCRs for each domain at 18 hpf, using the Signac framework. This analysis revealed that the most significantly enriched motifs are largely domain-specific (Figure 3D, Supplementary Dataset 4), and aligned with differentially expressed transcription factors in each domain. For instance, retinal DOCRs exhibited the highest enrichment for motifs associated to homeobox (LHX2, VSX1, VSX2, VAX2) and nuclear receptor (RORA, RORB, RORC) transcription factors, alongside the zinc-finger factor VEZF1 (Figure 3E).

### Identification and functional validation of cell-type specific cis-regulatory elements

To further investigate the regulatory logic underlying specification programs in the neural retina and adjacent forebrain domains, we sought to identify domain-specific cis-regulatory elements (CREs) from the multiomic dataset. To this end, DOCRs were integrated with cell-type-specific DEGs to highlight genomic loci potentially associated with active regulatory programs (see Methods). Candidate regions were further filtered based on their proximity to DEGs and the presence of binding motifs for TF-encoding genes differentially expressed in the same territory (Figure 4A and 4B). This integrative strategy established a coherent link between chromatin accessibility, TF activity, and gene expression.

**Figure 4.**
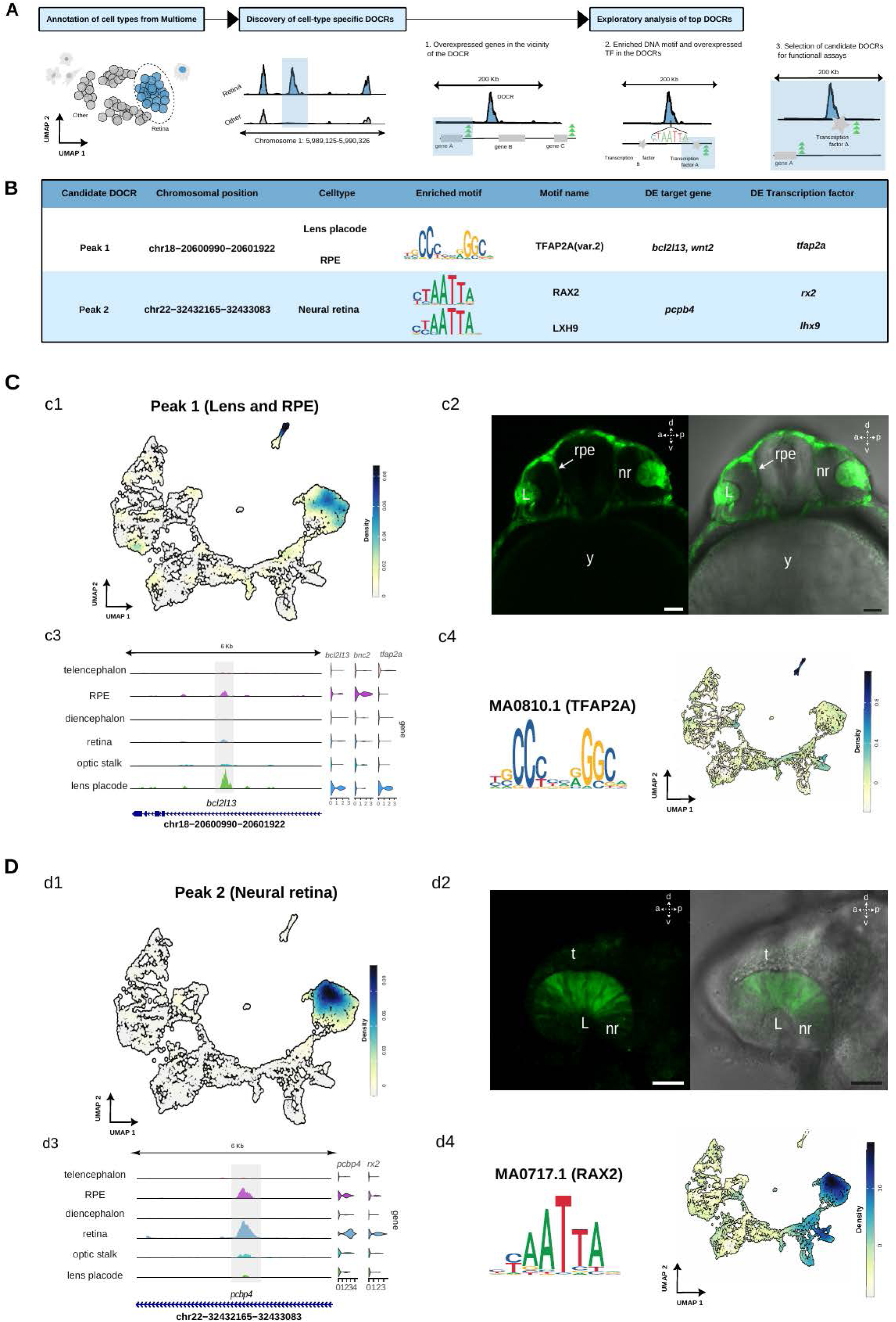
Identification and *in vivo* validation of domain-specific cis-regulatory elements. **(A)** Schematic overview of the cis-regulatory element (CRE) identification strategy. For each annotated domain in the 18 hpf multiome dataset, differentially open chromatin regions (DOCRs) were computed and intersected with nearby differentially expressed genes (DEGs). DOCRs containing enriched transcription-factor (TF) motifs, whose corresponding TFs were also differentially expressed within the same domain, were retained as candidate CREs. **(B)** Examples of candidate CREs prioritized through this pipeline. For each DOCR, genomic coordinates, enriched motif, associated DEG, and differentially expressed TF with a predicted binding site are shown. Two representative candidates are highlighted: a lens placode/RPE-associated element and a neural retina–specific element. **(C)** In vivo and multiomic validation of the RPE/lens candidate CRE. **(c1)** Density plot showing domain-specific chromatin accessibility of the DOCR across the 18 hpf multiome UMAP. **(c2)** Reporter assay using the ZED construct demonstrating specific GFP expression in the RPE and lens placode. **(c3)** Genomic view of the DOCR (chr18:20600890–20601932), located within the *bcl2l13* locus, with peaks showing higher accessibility in lens placode and RPE; violin plots display expression of nearby DEGs (*bcl2l13*, *bnc2*) and of *tfap2a*, encoding for a TF with a motif present in this region. **(c4)** *TFAP2A* motif logo (MA0810.1) and its motif-activity density across the multiome UMAP, strongest in the lens placode. **(D)** In vivo and multiomic validation of the neural retina candidate CRE. **(d1)** Density plot showing the DOCR’s accessibility pattern, enriched in neural retina. **(d2)** Reporter assay showing predominant expression in neural retina. **(d3)** Genomic view of the CRE (chr22:32432165–32433098) within the *pcbp4* locus, displaying increased accessibility in neural retina and RPE; violin plots show expression of *pcbp4,* as well as of *rx2*, encoding for a TF whose predicted motif is present in the region. **(d4)** RAX2 motif logo (MA0717.1) and motif-activity distribution, with strongest activity in neural retina and detectable activity across other anterior brain territories.

To assess the functional activity of these candidate CREs in vivo, a subset of the most statistically significant regions was cloned into ZED reporter constructs (Supplementary Figure S3A) and injected into zebrafish embryos at the single-cell stage. Almost all regions tested (8 out of 9) drove reporter expression in the predicted domains, validating the robustness and specificity of our approach. Representative examples of validated CREs are shown for the RPE/Lens (Figure 4C), neural retina (Figure 4D), dorsal diencephalon, telencephalon and optic stalk (Supplementary Figure S3B, 3C and 3D, respectively).

For the RPE/lens domains, the element chr18:20600890–20601932 exhibited increased chromatin accessibility, contained the binding motif of the enriched regulator TFAP2A (variant 2), and was located in proximity to the differentially expressed genes *bcl2l13* and *wnt2.* When tested *in vivo* using the ZED reporter assay, this region droved specific GFP expression in the RPE and lens placode, as predicted by the multiomic analysis (Figure 4C). Similarly, the candidate element chr22:32432165–32433098 displayed differential accessibility in the neural retina, was in the vicinity of the retina-enriched gene *pcbp4*, and contained predicted binding motifs for RAX2 and LHX9. The genes encoding for these transcription factors were also differentially expressed within the neural retina population, indicating a tightly connected regulatory logic. Consistent with these predictions, reporter assays confirmed specific activity of this element in the developing retina (Figure 4D). Candidate CREs in the remaining domains were likewise validated by reporter assays (Supplementary Figure S3B–D). The dorsal diencephalon element (chr5:29777849–29778766) contained DMBX and TCF7L motifs and was located near the differentially expressed gene *barhl1a*. In the telencephalon, the element chr22:16818184–16819067 harbored VAX2, EMX2, and DMRTA2 motifs and was positioned close to the telencephalic marker *nfia*. Finally, in the optic stalk, the region chr17:21346407–21347301 included ZIC2 motifs and was located near *vax1*, *eno4*, and *shtn1*. Notably, these three genes comprise a synexpression group, as the three of them displayed specific expression at the optic stalk.

To assess the extent to which the cis-regulatory architecture of the zebrafish eye field provides insight into its human counterpart, we examined the evolutionary conservation of the neural retina DOCRs we identified in zebrafish (Supplementary Figure S4). A total of 1,710 retinal DOCRs (log₂FC ≥ 1, adjusted p ≤ 0.05) were aligned to the human genome using *blastn*, retaining alignments of at least 50 bp. Of these regions, 1,100 showed no detectable conservation, whereas 610 DOCRs yielded significant sequence alignments in the human genome (2,100 hits in total). Among the conserved regions, 464 DOCRs (1,940 hits) mapped to loci not associated with orthologous gene pairs between zebrafish and human. In contrast, 136 DOCRs (160 hits) were located near human loci whose orthologous genes were also proximal to the corresponding DOCRs in zebrafish. These 136 conserved DOCRs were linked to 170 human orthologous genes (Supplementary Dataset 5). Notably, some of these genes encode key transcription factors implicated in retinal patterning, including *PAX6*, *RAX*, *SIX3*, *VAX1*, *VSX2*, and *OTX1*, as well as signaling regulators such as *ALDH1A3*, *SMOC1*, and *FZD5*. Importantly, all of these genes have been previously implicated in human microphthalmia, anophthalmia, and coloboma (MAC) syndromes, underscoring the relevance of conserved cis-regulatory elements identified in zebrafish for understanding human ocular development and disease.

### Multi-omics analysis of gene regulatory networks across eye and forebrain domains at 18 hpf

To systematically investigate the regulatory programs underlying eye and anterior forebrain specification in zebrafish, we integrated our single-nucleus RNA-seq and ATAC-seq data from 18hpf using MatchaiRen software framework *(Multi-omics analysis of transcriptomics and chromatin accessibility in Regulatory network inference)* (Figure 5A). This approach enabled the reconstruction of putative, territory-specific regulatory networks by linking DEGs to differentially expressed transcription factors (DETFs) via nearby differentially open chromatin regions (DOCRs) containing enriched TF binding motifs (See methods). For each inferred network, we quantified node connectivity and applied a weighted scoring function to rank highly connected regulatory hubs across all territories (Figure 5B, Supplementary Figure S5-S6, Supplementary Dataset 6).

**Figure 5.**
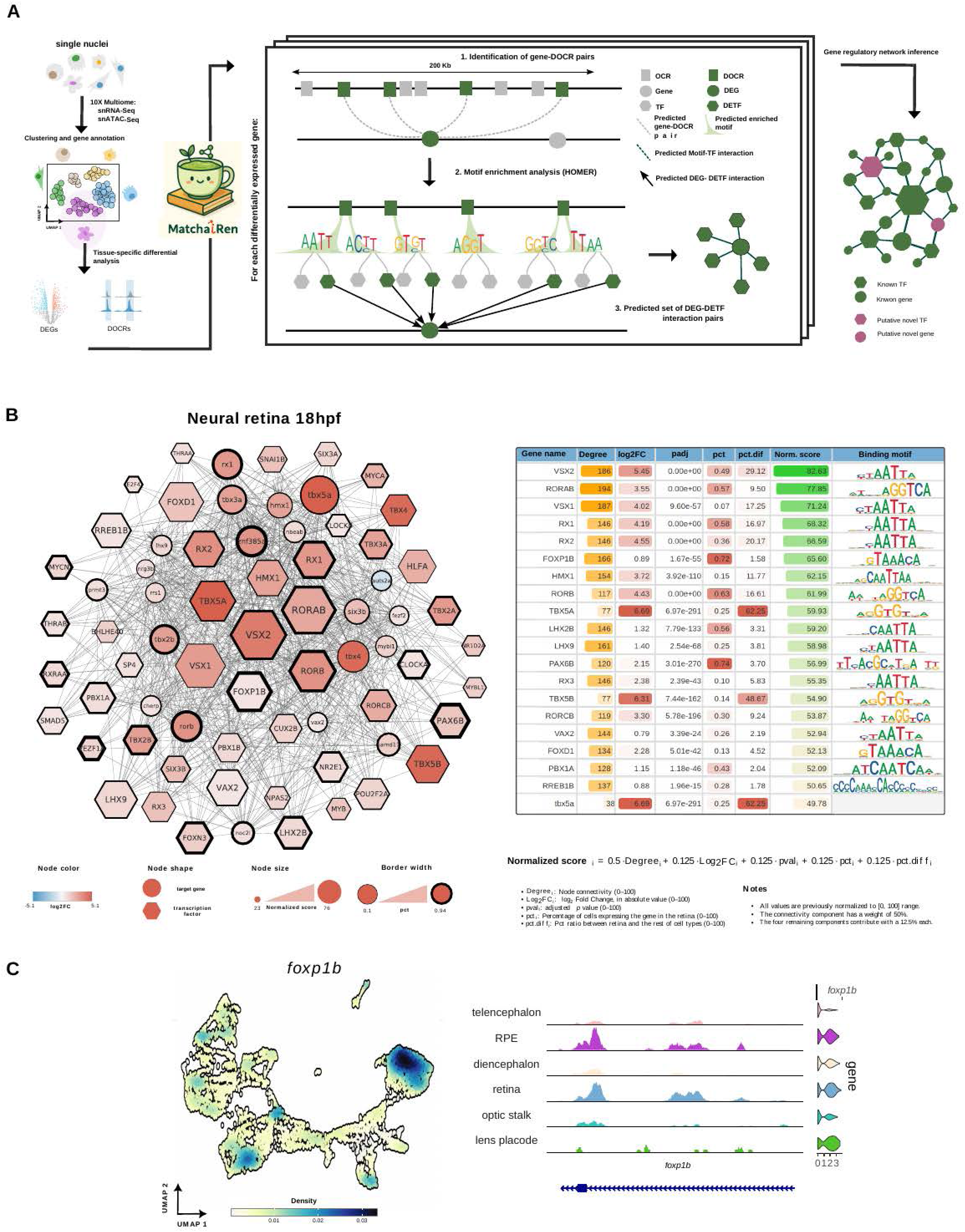
Gene regulatory network inferred from multiome data reveals highly connected hubs at the retinal domain. **(A)** Schematic representation of the GRN inference workflow (MatchaiRen). For each annotated domain, DEGs and DOCRs from the multiome dataset are integrated to identify candidate TF–target interactions. DOCRs are linked to nearby DEGs; enriched motifs are identified within each DOCR, and interactions are retained when the corresponding TF is also differentially expressed in the same domain. The aggregation of all DOCR-supported TF–gene pairs yields a domain-specific gene regulatory network. **(B)** Inferred gene regulatory network for the neural retina domain. Highly connected nodes correspond to known regulators of retinal identity, including *vsx2*, *vsx1*, *rorab*, *rx1/2*, *lhx9* and *lhx2b*. A ranked table reports the weighted node score (normalized_score), integrating changes in gene expression, cellular prevalence, and connectivity. This ranking highlights canonical retinal regulators but also identifies less characterized candidates such as *foxp1b*. **(C)** Expression and chromatin-accessibility patterns of *foxp1b*. A density UMAP shows its spatial expression enriched in the neural retina cluster. Genomic tracks reveal domain-specific chromatin accessibility at the *foxp1b* locus, supporting its predicted role as a regulatory hub in early retinal development.

Focusing on the retina, the inferred network recapitulated many transcription factors previously described as central regulators of neural retina specification (*6*). Among these, *vsx2*, *rorab* and *vsx1* exhibited the highest normalized scores and connectivity degrees, followed by *rorb*, *rx1/2, foxp1b* and members of the LHX family, including *lhx9* and *lxh2b* (Figure 5B). Because the weighted scoring integrates orthogonal layers of information—such as gene expression, gene prevalence across cells, and network connectivity—the resulting hierarchy of nodes not only highlights highly connected regulators but also prioritizes those consistently enriched across the retinal domain. This provides a robust and integrated representation of the transcriptional landscape governing neural retina specification.

### Foxp1b as a conserved regulatory hub in retinal development

Among the highly connected TFs identified in the inferred retinal network, *foxp1b*, a gene with a broad expression in the central nervous system but poorly characterized in the retina (*41*), emerged as a differentially expressed central node. Its enrichment at the retinal domain, as well as its high normalized score and connectivity degree within the neural retina network, underscores its potential role as a key regulator of retinal specification (Figure 5 B, C).

Notably, *foxp1b* is located in chromosome 6 in close genomic proximity to *mitfa,* a major regulator of RPE specification. This genomic arrangement is conserved in humans, where *FOXP1* and *MITF* are located at 3p13–p14), comprising a deeply conserved syntenic block maintained across vertebrates, including tetrapods (human, mouse), holosteans (spotted gar), and teleosts (zebrafish, medaka, stickleback), despite the whole-genome duplication event (Supplementary Figure S7A). The conserved proximity of these genes suggests selective pressure to maintain the architecture of the locus, potentially reflecting a coordinated regulatory control during eye development. Despite this genomic linkage, *foxp1b* and *mitfa* exhibit mutually exclusive expression patterns in the retina and RPE domains, respectively, as revealed by our single-cell analyses and confirmed by HCR experiments (Supplementary Dataset 2, Supplementary Figure S7B). To further investigate the regulatory logic of the locus, we examined available Micro-C data in human ESCs (*42*), together with zebrafish chromatin accessibility profiles from neural retina and RPE populations (*13*). Human Micro-C data indicated that *FOXP1* and *MITF* reside in neighboring Topological Associating Domains (TADs), separated by a strong insulating boundary (Supplementary Figure S7C). Consistently, bulk ATAC-seq analysis in zebrafish from isolated neural retina and RPE populations at 23hpf (*13*) revealed a partitioned regulatory landscape, with RPE-specific CREs associated to *mitfa* and neural retina-specific CREs linked to *foxp1b* (Supplementary Figure S7C). Together, these observations suggest that *Foxp1* and *Mitf* form a bipartite eye regulon with a conserved regulatory architecture across vertebrate species.

Taking advantage of stablished and highly efficient CRISPR-Cas9 methods for the generation of small deletions at F0 in the zebrafish genome (*43*), we functionally validated the role *foxp1b* in retinal specification in both zebrafish and medaka embryos (Figure 6). Injection of two independent sgRNA pairs targeting exons 1 and 6 of the zebrafish *foxp1b* gene (Figure 6A), resulted in a significant reduction of the eye size in 18hpf crispant embryos compared with control siblings, with an average area reduction of 21-29% (Figure 6B-C). Similarly, injection of two independent sgRNA pairs targeting exons 1 and 3/4 of the medaka *foxp1b* gene (Figure 6D) led to a marked reduction in eye size in 38hpf crispant embryos, with an average area reduction of 43–68% relative to controls (Figure 6E-F). Notably, a small proportion of the embryos exhibited severe unilateral microphthalmia. Together, these results indicate a conserved requirement for *foxp1b* during retinal development in teleosts.

**Figure 6.**
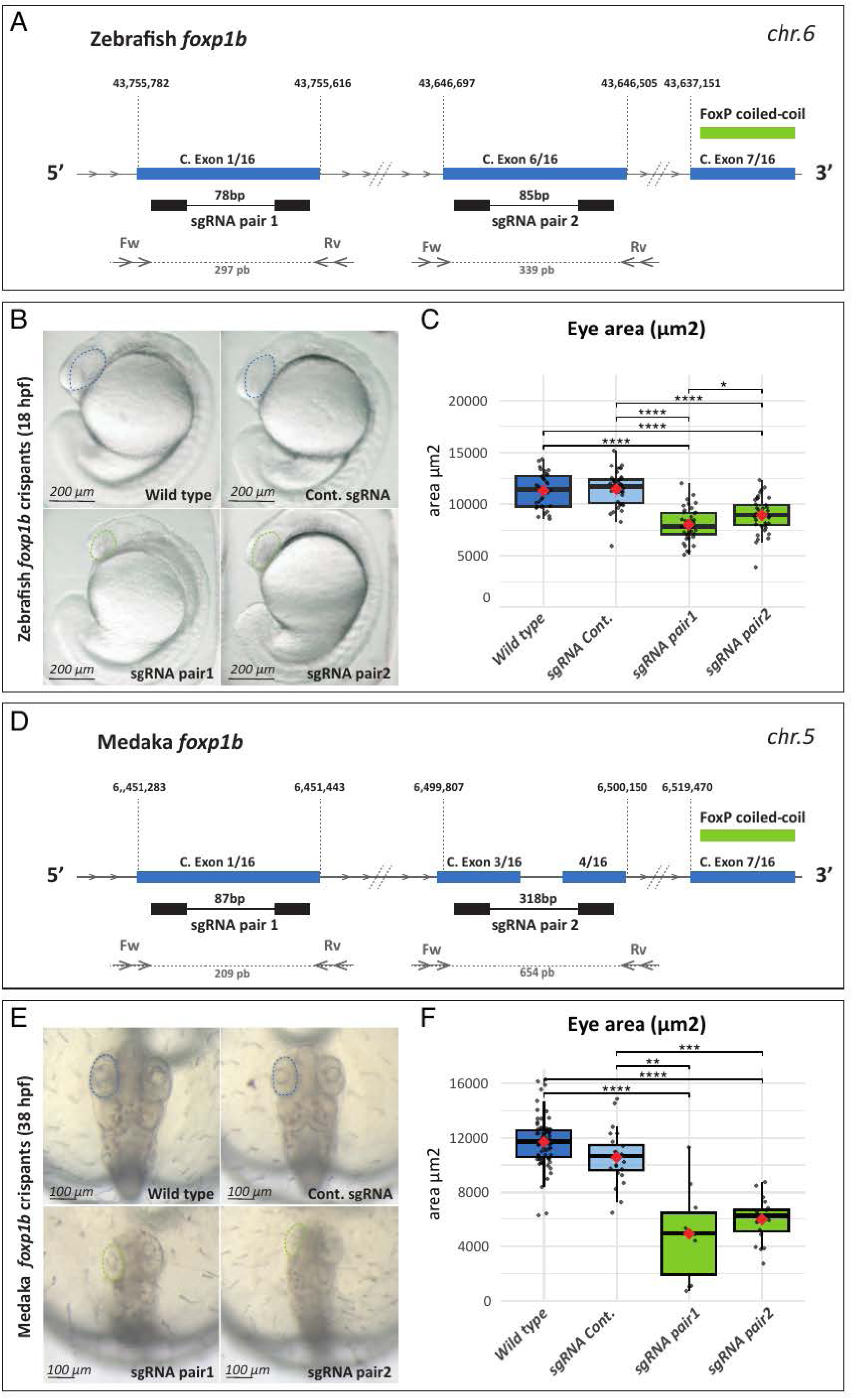
CRISPR-Cas9 editing of the *foxp1b* locus in zebrafish and medaka. (A) Schematic representation of exons 1, 6 and 7 (blue boxes) corresponding to *foxp1b* in the zebrafish genome at chromosome 6; genomic coordinates are indicated for each exon. Black boxes indicate the position of specific sgRNAs pairs and their relative distance shown in each case. Primers for mutagenic assessment are depicted as opposing arrowheads. (B) Lateral view of 18hpf zebrafish embryos showing area of the optic primordium for wild type and embryos injected with a control sgRNA pair (medaka *efs*; dashed blue line), as well as injected with *foxp1b* sgRNAs pairs (dashed green line). (C) Quantification of eye area (µm2) in control (blue) and crispant (green) zebrafish embryos. Black dots represent individual optic vesicle measurements. Data are the mean ± SD (****p<0.000001; *p<0.05). (D) Schematic representation of exons 1, 3, 4 and 7 (blue boxes) corresponding to *foxp1b* in the medaka genome at chromosome 5; genomic coordinates are indicated for each exon. Black boxes indicate the position of specific sgRNAs pairs and their relative distance shown in each case. Primers for mutagenic assessment are depicted as opposing arrowheads. (E) Dorsal view of 38hpf medaka embryos showing the area of the optic cup for wild type and embryos injected with a control sgRNA pair (medaka *efs*; dashed blue line), as well as injected with *foxp1b* sgRNAs pairs (dashed green line). (F) Quantification of eye area (µm2) in control (blue) and crispant (green) medaka embryos. Black dots represent individual optic vesicle measurements. Data are the mean ± SD (****p<0.00001; ***p<0.0001; **p<0.001).

To further investigate the molecular consequences of *foxp1b* loss of function in zebrafish, we compared the transcriptomes of 18hpf *foxp1b* crispants and wild type siblings using bulk RNAseq. This analysis identified a total of 1,138 significantly dysregulated genes in crispant embryos ([log₂FC] ≥ 0.5; adjusted p-value ≤ 0.05), including 618 upregulated and 520 downregulated genes (Supplementary Figure S8A; Supplementary Dataset 7). Gene ontology analysis of the downregulated genes revealed “optic vesicle” as one of the most significantly enriched terms (Supplementary Figure S8B; Supplementary Dataset 7). Moreover, many of the genes predicted by our MatchaiRen analysis to be both potential *foxp1b* targets and central nodes of the retinal network (Figure 5) appeared downregulated in *foxp1b* crispants (Supplementary Figure S8C; Supplementary Dataset 7). Collectively, these findings support a central role for *foxp1* as a key regulator for the specification of the neural retina domain.

## Discussion

Using a combination of scRNAseq and snMultiome profiling, we dissected the architecture of the gene regulatory networks that sustain the identity of each of the major compartments of the developing zebrafish eye. This work offers a reference dataset that will contribute to both understanding the regulatory logic underlying vertebrate anterior brain patterning and gaining insight into the aetiology of common hereditary eye malformations, such as MAC syndromes. Our analyses focus on a precise developmental window, after compartment identities are established but before the onset of neuronal differentiation, and are extended to neighbouring forebrain territories. This study complements previous single-cell multiome and chromatin accessibility studies performed at later stages, which primarily focused on retinal differentiation and neuronal fate acquisition. Such studies, conducted in representative vertebrate species, including human, mouse, chicken, zebrafish, and lamprey (*36–38, 44, 45*), have revealed a deeply conserved cis-regulatory code governing neuronal subtypes specification, despite extensive evolutionary turnover of individual regulatory elements (*44*). These findings are in agreement with the well conserved histological organization and transcriptomic profiles of the vertebrate eye (*5*). Our datasets at earlier stages are also in line with a broad conservation of the phylotypic vertebrate brain (*46*). Indeed, most of the highly interconnected nodes identified in each territory-specific regulatory network correspond to transcription factors previously described as central regulators in other vertebrates and implicated in hereditary eye malformations in humans (*9*).

Within the framework of a broadly conserved anterior brain blueprint, our study provides insight into the relationships between transcriptional programs operating across distinct domains. Previous work from our laboratory demonstrated that the acquisition of the RPE fate requires the repression of the neural retina program (*13*). Extending this model, our RNA velocity analyses, now integrating information from the eye and adjacent forebrain territories, suggests that the proliferative neural retina program may also represent a naive transcriptional state for the optic stalk and telencephalic tissue, and thus likely for all most anterior neural derivatives (Figure 2). Importantly, this conclusion does not imply a lineage relationship between the different neural domains, which cannot be inferred from RNA velocity studies in the absence of complementary lineage tracing approaches. Rather, our data support the existence of a shared transcriptional program among anterior brain derivatives that closely resembles that of the neural retina compartment. This interpretation is in agreement with genetic evidence showing that the acquisition of the optic stalk and telencephalic identities likewise depends on the repression of the neural retina program (*47–49*).

The observation that most of the central nodes within the eye-specific regulatory networks we identified were already known from human genetic studies suggests that discovery of novel core components is approaching saturation. This constrains potential explanations for the missing heritability of MAC syndromes, shifting attention towards complex genetic interactions and non-coding variants as likely causal mechanisms.

In particular, non-coding variants affecting ultra-conserved CREs may account for substantial fraction of this missing heritability. Ultra-conserved elements are exceptionally depleted of common variants in human populations (i.e. at levels comparable to missense variants in protein-coding regions), reflecting strong purifying selection (*50*). Although a limited number of non-coding pathogenic variants have already been reported in MAC patients (*12, 51*), systematic efforts to interrogate CREs in human patients with rare eye diseases have only recently begun (*52*). Our study offers a reference framework for the inference of domain-specific CREs (Supplementary Dataset 3) and includes the validation of most significant regions using transgenesis assays (Figure 4, Supplementary Figure S3). To assess the translational relevance of our findings to human disease, we analysed 1710 retinal-specific zebrafish regulatory elements and identify human ultra-conserved CREs associated with 170 orthologous loci. These loci are significantly enriched in genes encoding TFs previously implicated in MAC syndromes (Supplementary Figure S4). Given that synteny-based algorithms can reliably detect functionally conserved CREs across distantly related vertebrate species, even beyond sequence-conservation (*53*), our datasets are likely to provide valuable guidance for prioritizing candidate non-coding regions to be surveyed in genetically unresolved MAC families.

To integrate our single-nuclei multiomic data, we developed MatchaiRen, a gene regulatory network inference tool designed to reconstruct regulatory programs in spatially defined developmental domains. A growing number of methods have been developed to infer gene regulatory networks from single-cell datasets (e.g., ANANSE, SCENIC+, CellOracle), often integrating chromatin accessibility, transcription factor motif information and curated regulatory annotations (*34*). While these approaches are powerful and widely used, their application to developmental systems poses specific challenges. In particular, transcription factor–motif associations and regulatory annotations are more complete in mammals, whereas non-mammalian model organisms often rely on sparse or less curated resources. In addition, many existing frameworks tend to generate large and densely connected networks, which can be difficult to interpret and often require extensive downstream analyses. Rather than relying on predefined regulatory annotations, MatchaiRen adopts a comparison-based strategy, in which differential activity is used as a structural constraint, focusing on genes and regulatory elements that are specifically active within a given domain. This design facilitates the refinement of hierarchical interactions but also imposes some limitations. In particular, TF with very broad tissue expression tend to be underweighted. This was the case when we analyzed the neural retina network for a couple of neural TF-encoding genes, *sox2* and *meis1*, which despite their known role in retinal specification (*54, 55*) failed to be recovered as central nodes in our analysis. Importantly, MatchaiRen does not aim to reconstruct global regulatory networks. Rather, it leverages contrast across domains to identify the subset of regulatory interactions most likely to define domain identity or functional state. By emphasizing domain-specific regulatory programs over global connectivity, MatchaiRen yields more compact and interpretable networks. When applied to early zebrafish development, the framework successfully recovered known regulators and identified new candidate factors involved in eye and forebrain patterning.

To validate the analytical power of our approach, we focused on the well-known retinal regulatory network and investigated the function of *foxp1b,* whose role in retinal specification was poorly characterized despite its predicted high connectivity within the network. We show that *Foxp1* and *Mitf* form a bipartite eye regulon that is conserved across vertebrates; and we functionally validated a role for *foxp1b* in retinal specification in both medaka and zebrafish. In mice, constitutive targeting of *Foxp1* results in early embryo lethality due to severe cardiac defects (*56*). Although this study hinted to a mild microphthalmia in *Foxp1* -/- embryos, these observations were not further confirmed in retina-specific conditional models. Despite *Foxp1* is expressed in early retinal precursors in mice, no overt early eye defects have been described following conditional inactivation using either *Dkk3-Cre* (*57*) or *Six3-Cre* drivers (*58*). This contrasts with the microphthalmic phenotype we observed in *foxp1b* teleost crispants. This discrepancy may reflect a differential regulatory influence of *Foxp1* in different vertebrate species. Alternatively, the *Cre* divers used in these studies were selected to interfere with retinal differentiation rather than early tissue specification, and therefore an incomplete early inactivation of the gene cannot be ruled out. Interestingly, *Foxp1* has proved essential to maintain the identity of early retinal precursors and to regulate their competence to generate early-born neuronal types (*58*), thus supporting a primary role in the mice neural retina GRN. In humans, heterozygous *FOXP1* mutations cause a dominant neurodevelopmental disorder known as FOXP1 syndrome, characterized by intellectual disability, language delays and autism spectrum disorder (*59*). Remarkably, severe eye abnormalities have also been reported in a subset of affected individuals (≈5%,), including coloboma (*60, 61*), microphthalmia, and even unilateral anophthalmia (*62, 63*). Taken together, these reports support a role for *FOXP1* in structural eye phenotypes in human patients and are in line with a conserved role for *Foxp1* genes in neural retina specification across vertebrates. Although our validation efforts primarily focused on the neural retina domain, eye malformations can arise from defects in the formation of any optic cup components. Accordingly, our analyses identifying domain-specific CREs and highly connected nodes within each of the eye-specific regulatory networks provide a reference framework that may guide future investigations in genetically undiagnosed MAC patients.

## Materials and Methods

### Experimental model and subject details

All experiments performed in this work comply European Community standards for the use of animals in experimentation and were approved by the Pablo de Olavide University and Junta de Andalucía ethical committees (license #02/11/2021/165). Zebrafish AB/Tübingen (AB/TU) and medaka iCab wild-type strains were staged, maintained and bred under standard conditions (*64, 65*). The zebrafish line used in flow cytometry experiments, *Tg(rx3:Gal4;UAS:GFP),* previously described (*39, 66*), was also maintained under standard conditions.

### Embryo dissociation and single-cell libraries preparation

*For scRNAseq experiments:* Approximately 500 whole embryos, either wild type or *Tg(rx3:Gal4;UAS:GFP)*, were processed for each developmental stage: i.e. 15, 18 and 23 hpf. Cell dispersions were obtained as previously reported (*67*). Briefly, embryos were first dechorionated for 10 min with pronase (30 mg/ml in 0.3X Danieau’s solution) and, after enzyme quenching (10 % FBS in Danieau 0.5X), embryos were deyolked on ice as reported (*67*). Deyolked material was pelleted and cell dissociated by 10 min incubation in 3 ml of FACSMax solution (Amsbio, T200100) on ice. Dissociated cells were resuspended in 1 ml 2% FBS/HBSS and filtered 3x through a 70 μm cell strainer before sorting. Cell suspensions were sorted using a SONY MA900 (laser 488/561). 7-AAD (7-amino actinomycin D) was used to discriminate dead cells. Relaxed gating parameters were applied to recover cells expressing a broad range of GFP levels (Supplementary Dataset 8). Recovered cells were resuspended in 2% FBS/HBSS and cell number (450-900 cells/μl) and viability assessed using TC-20 cell counter (Bio-Rad) after trypan blue staining. Once cell viability was confirmed to be over 90%, 20.000 cells were loaded onto a Chromium Controller, G chip (10× Genomics, Pleasanton, CA, USA). For the generation of the scRNAseq libraries, we used a Chromium Next GEM Single Cell 3ʹ Kit v3.1, according to the manufacturer’s instructions. Sequencing was performed on a DNBSEQ-T7 system (BGI Group, Shenzhen, China). Sequencing depth was 43,792, 39,184, and 94,027 reads/cell at 15, 18 and 23 hpf respectively.

*For snMultiome experiments:* 150 dissected embryo heads were harvested and cell suspensions obtained as described in the previous section. For nuclei isolation. we followed the 10X Genomics protocol for single-cell multiome sequencing (CG000338). Briefly, dispersed cells were lysed for 75 sec by gentle pipetting mix on chilled lysis buffer. Washed buffer was then added to stop outer membranes lysis and material was filtered (70 µm nylon filters). Nuclei were cleaned 2x in washed buffer, harvested by centrifugation 5min 300g at 4°C and further filtered (40 µm nylon filters). Recovered nuclei were resuspended in diluted nuclei buffer (CG000338) and nuclei number estimated using TC-20 cell counter (Bio-Rad). Nuclear viability was assessed in parallel after DAPI staining using a Neubauer Chamber. 14,582 nuclei were loaded onto a Chromium Controller, J chip (10× Genomics, Pleasanton, CA, USA), and processed using the Chromium Next GEM Single Multiome ATAC + Gene expression Kit, according to the manufacturer’s instructions. Sequencing was performed on a DNBSEQ-T7 system (BGI Group, Shenzhen, China). Sequencing depth was 33,880 reads/cells (RNA-seq) and 28,915 reads/cells (ATAC-seq).

### CRISPR genome editing and crispants analysis

To interfere with *foxp1b* function, two independent single guide RNAs (sgRNAs) pairs targeting the corresponding loci in the zebrafish and medaka genomes (Figure 5) were designed using CRISPRscan (*68*). Primers for sgRNA generation (Supplementary Dataset 9) were aligned by PCR to a universal CRISPR primer and the resulting product further purified and used as a template for sgRNA synthesis by in-vitro transcription (*69*). To target individual loci, a solution containing the specific *foxp1b* sgRNAs mix (80 pg), SpCas9 protein (250 pg), and GFP mRNA (25 pg) as a tracer, were injected into one-cell-stage zebrafish and medaka embryos. Control embryos were injected with an irrelevant sgRNA mix directed against the medaka gene *efs*, which yield no overt phenotype. Oligos used for screening the presence of genomic DNA deletions are detailed in (Supplementary Dataset 9).

For crispants’ phenotypic analysis, embryos were placed on 1.2% agarose lanes and eye areas measured using FIJI/ImageJ (Version 1.50i). Quantitative data were evaluated using R Studio software (2023.06.0+421 version). A one-way repeated-measures analysis of variance (ANOVA) followed by the Holm-Sidak test for pairwise comparison was used on normally distributed samples, while Kruskal-Wallis test followed by Dunn’s test with Bonferroni correction was used on not normally distributed samples (Shapiro-Wilk and Levene tests).

### Bulk RNAseq experiments

Total RNA was extracted from 18hpf WT and *foxp1b* crispant embryos using the easy-BLUE Total RNA Extraction Kit (Intron Biotechnology, Seongnam, Korea), following the manufacturer’s protocol. Genomic DNA contamination was removed by treating RNA samples with the TURBO DNA-free Kit (Ambion, #AM1907). Three biological replicates, each consisting of 12 embryos, were included for each condition analyzed. RNA-seq libraries were prepared and sequenced by BGI using the DNBseq platform, generating paired-end reads of 100 bp in length, with a minimum of 30 million reads per sample.

### Functional analysis of cis-regulatory elements by transgenesis

Selected putative cis-regulatory regions were PCR amplified from gDNA using primers designed to span the open chromatin regions (Supplementary Dataset 9). Corresponding bands were gel purified and cloned into a pCR™8/GW/TOPO® vector (Thermo Fisher Scientific) for bacterial amplification. Once purified and verified, plasmids were recombined into the Tol2-compatible vector Zebrafish Enhancer Detection (ZED) (*70*). Final constructs were confirmed by PCR, for the presence of the putative non-coding regulatory sequences, and injected into one-cell zebrafish embryos following the described protocol (*70*). F0 transgenic embryos were screened from 15 to 24 hpf in an Olympus SZX12 stereo fluorescence microscope. Positive embryos were selected for further growth to adulthood to obtain stable transgenic lines.

### HCR

HCR RNA-FISH technique was performed following the protocol by Molecular Instruments (*71*) adapted to whole-mount zebrafish embryos. 18hpf zebrafish embryos were fixed in 4% formaldehyde overnight at 4°C and washed with PBS to stop the fixation. Subsequently, embryos were dehydrated through an increasing series of methanol (MeOH) solutions in PBS, for their storage in 100% MeOH at -20°C. Before use, embryos were rehydrated in an inverse series of MeOH/PBST. Permeabilization with Proteinase K was skipped. Then, they were incubated in a pre-hybridization buffer at 37°C, followed by the addition of specific pairs of DNA probes (28 against *foxp1b* and 19 against *mitfa*) and overnight incubation at the same temperature. Buffers and hairpins were ordered to Molecular Instruments, Inc. DNA probes were designed as described (*72*) and ordered to IDT as Oligo-Pools. Washes were performed to remove unbound probes, and the fluorescent signal was triggered and amplified using fluorophore-associated pair of hairpins. Finally, embryos were washed to remove the excess of hairpins and stained with DAPI to be visualized in a confocal microscope Leica Stellaris5.

### Single-cell RNA-seq preprocessing and integration across developmental stages

Single-cell RNA-seq data were generated from zebrafish embryos at 15, 18, and 23 hours post-fertilization (hpf). Raw sequencing reads were aligned and quantified using *10X Genomics Cell Ranger* v6.1.2 (*73*) against the *Danio rerio* reference genome from *NCBI* (*GRCz11*). Low-quality cells and cells with excessive mitochondrial RNA content were removed. Expression matrices were normalized, log-transformed, and integrated across developmental stages using *Seurat* v5.0.3 (*74*) with reciprocal PCA-based anchoring, enabling downstream comparison while preserving stage-specific identities. Dimensionality reduction was performed using PCA followed by UMAP, and clustering was carried out with the Louvain algorithm testing multiple resolution parameters to refine cluster granularity. Cell type annotation was performed using a marker-based enrichment strategy: differentially expressed genes (DEGs) from each cluster were compared to curated markers from ZFIN (*20*) to assign cell identities consistently across developmental stages.

### Cell annotation

To assign biological identities to transcriptional clusters, we used a marker-based enrichment strategy. For each cluster, marker genes were computed using the *FindMarkers* function from the *Seurat* package, using the *Wilcoxon* rank-sum test and retaining positive markers with adjusted *p* ≤ 0.05 and log₂FC threshold ≥ 0.5. These genes were compared against a relational anatomical annotation database from ZFIN, which maps genes to specific anatomical terms. The statistical significance of enrichment for each anatomical term was calculated using a hypergeometric test: let *N* be the total number of detected genes in the experiment, *K* the number of genes associated with a given anatomical term, *n* the number of cluster-specific overexpressed genes, and *k* the number of cluster genes associated with that term. The probability of observing *k* or more genes by chance was computed as:

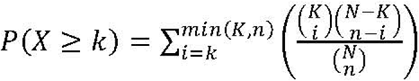

P-values were adjusted for multiple testing using the Benjamini-Hochberg method. The anatomical term with the lowest adjusted p-value was assigned as the primary annotation for the cluster.

### Trajectory inference and Dynamic Gene Expression Analysis

To explore differentiation trajectories, we performed RNA velocity and latent time inference analyses using *scVelo* v0.2.5 (*75*). For the neural retina domain, subsets of cells annotated as neural retina and proliferative region were integrated and combined with spliced/unspliced transcript counts from *Velocyto*-generated v0.17.17 loom files (*76*). After filtering, normalization, and moment computation on the neighborhood graph, the dynamical model was applied to estimate RNA velocities and latent time. Genes were ranked according to multiple dynamical parameters (likelihood, alpha, beta, gamma, variance), and the top candidates from each ranking were intersected to obtain a robust velocity-informed gene set. This set was further intersected with domain-specific DEGs. Expression dynamics of these intersected genes were visualized in heatmaps, with cells ordered along latent time and genes ordered by their peak expression timing, revealing early- versus late-activated gene programs.

### Single-nuclei multiome (RNA + ATAC) analysis

Single-nuclei multiome data were generated from pooled 18 hpf embryos using the *10x Genomics Multiome* platform. Raw data were processed with *Cell Ranger ARC* v2.0.2 (*77*) to obtain paired RNA and ATAC count matrices. Accessible chromatin peaks were directly called by Cell Ranger ARC, which provides peaks as ATAC features. Quality control included filtering nuclei based on RNA and ATAC features (e.g., number of detected genes, fragments in peaks, transcription start site enrichment, nucleosome signal) and potential doublets were identified and removed at this stage using *scDblFinder* v1.16.0 (*78*). Joint dimensionality reduction was performed with weighted nearest neighbors (*WNN*) in *Seurat* and *Signac* v1.13.0 (*79*), integrating RNA and ATAC modalities. Clusters were defined from the multimodal graph and annotated using the same marker-enrichment strategy described for scRNA-seq, ensuring consistent cell type assignment across RNA-seq and multiome datasets.

### Differential expression and chromatin accessibility analysis

For each annotated cell type, differentially expressed genes (DEGs) were identified using the *FindAllMarkers* function in *Seurat*, with thresholds of log fold-change ≥ 0.25 and minimum detection in 5% of cells, controlling for technical covariates where appropriate. Similarly, for each annotated cell type, differentially open chromatin regions (DOCRs) were identified using *FindAllMarkers* on the ATAC assay in *Signac*, employing a logistic regression test (test.use = “*LR*”), controlling for sequencing depth and technical covariates.

### Motif enrichment analysis

To identify transcription factor (TF) motifs associated with domain-specific regulatory programs, we performed motif enrichment analysis using the *Signac* R package. Motif position frequency matrices (PFMs) were retrieved from the *JASPAR 2020 CORE* vertebrate collection (*80*) and incorporated into the multiome Seurat object via the *AddMotifs* function. For each of the six major domains (neural retina, RPE, lens placode, optic stalk, telencephalon, and diencephalon), DOCRs were defined relative to all other cell types based on adjusted *p* ≤ 0.05, log₂FC ≥ 1, and a minimum accessibility ratio of 1.5. Domain-specific DOCRs were matched to a background set of peaks with similar GC content and accessibility using *MatchRegionStats*. Enriched motifs were then identified with *FindMotifs*, and motif names were standardized to remove redundant variant annotations. To associate enriched motifs with putative regulators active in each domain, we cross-referenced the human motif gene names with zebrafish orthologs (ZFIN–Ensembl mapping) and retained only those corresponding to transcription factors significantly upregulated in the same domain (*adjusted p* ≤ 0.05; log₂FC ≥ 1).

### Identification of putative CREs

DOCRs were first intersected with cell-type-specific DEGs: for each DOCR, we searched a ±100 kb window around its genomic coordinates to identify DEGs nearby that could represent potential regulatory targets. Regions satisfying this proximity criterion were further analyzed by motif enrichment analysis using *HOMER2* v5 (*81*), with vertebrate transcription factor motifs obtained from the *JASPAR* database. Candidate regulatory regions were prioritized when enriched motifs corresponded to transcription factors (TFs) whose genes were also differentially expressed in the same cell type, providing a consistent link between chromatin accessibility, TF expression, and putative target gene regulation.

### Conservation analysis

To investigate evolutionary conservation of DOCRs identified in retinal cells, we selected candidates with log2FC ≥ 1 and adjusted p-value ≤ 0.05. These regions were aligned to the human genome using *blastn* v2.9.0+ (*82*), and alignments with a minimum length of 50 bp were retained. To associate each hit with nearby genes, we expanded zebrafish DOCR coordinates to ±100 kb and human hit coordinates ±200 kb, and intersected these regions with annotated genes from each species. Orthologous relationships between zebrafish and human genes were then used to identify DOCRs with conserved gene neighborhoods. This analysis yielded a subset of DOCRs that are evolutionarily conserved and associated with orthologous genes, providing a framework for downstream functional annotation and integration with transcription factor networks. Detailed scripts and parameters for sequence alignment, gene annotation, and ortholog matching are provided as supplementary code on GitHub.

### Gene regulatory network inference with MatchaiRen

To reconstruct cell-type-specific regulatory networks, we developed *MatchaiRen v0.1,* a Snakemake-based pipeline for the integrative analysis of transcriptomic and chromatin accessibility data. The pipeline was implemented using *Snakemake v9.6.2*; (*83*) and executed within a *Docker* container to ensure reproducibility. For each cell type, putative target genes were defined as DEGs. Candidate regulatory regions were defined as DOCRs located within ±100 kb of the gene body. These regions were scanned for transcription factor binding motifs using *HOMER*, with the *JASPAR* vertebrate motif database as reference. To enable motif-based inference in zebrafish, *JASPAR* human and mouse motifs were mapped to orthologous transcription factors via gene orthology from ZFIN.

Transcription factors that were both upregulated in a given cell type and enriched for binding motifs in nearby DOCRs were considered candidate regulators. The resulting TF–target relationships were used to construct cell-type-specific bipartite regulatory graphs, providing a mechanistic framework of transcriptional control during early eye development. For each network, we calculated node connectivity and derived a weighted scoring function to rank highly connected hubs across all territories.

This scoring function integrates several orthogonal parameters beyond connectivity, including the magnitude and statistical significance of differential expression (*log₂FC* and adjusted *p*-value), the proportion of cells in which each gene is detected, and the relative enrichment between the target domain and the rest of the domains. In this way, the score reflects not only the topological centrality of each node, but also its transcriptional prominence and domain specificity, reducing bias toward highly connected but lowly expressed genes. By combining these metrics, the scoring function highlights genes that are both strongly network-integrated and biologically relevant.

### Bulk RNAseq computational analysis

Reads were aligned to the *danRer11* zebrafish genome using *STAR* v2.7.10b (*84*). The transcript abundance was quantified with *featureCounts* v2.0.6 (*85*) and differential gene expression analysis were performed using *DESeq2* v1.49.2 (*86*), applying an adjusted p-value threshold of ≤ 0.05. To interpret differentially expressed genes, we performed a GO enrichment analysis adapted from our cell annotation strategy. Gene sets (up- and downregulated) were tested for enrichment of ZFIN anatomical terms using a hypergeometric test with *Benjamini–Hochberg* correction, and the resulting terms are reported ranked by adjusted p-value.

## Data and code availability

Single-cell RNAseq, single-nuclei multiome, and bulk RNAseq datasets are available in the Gene Expression Omnibus (GEO) repository (https://www.ncbi.nlm.nih.gov/geo) under the following accession numbers (GSE316738; GSE316739, GSE318207): Custom analysis scripts used in this study are available at https://github.com/rendon92/Macho_paper_2026. The MatchaiRen pipeline used for gene regulatory network inference is available at https://github.com/rendon92/MatchaiRen_v0.1.

## Acknowledgements

We thank Corín Díaz Ramos and Katherina García from the Advanced Light Microscopy and Citometry facilities at the CABD for their excellent technical assistance. We thank Rodrigo Young, Javier López-Ríos; Gretel Nusspaumer, and Ana Burgos for the scientific discussions and critical input on the manuscript. This work was supported by grants awarded to JRMM from the Spanish Ministry of Science, Innovation and Universities (AEI) (references PID2023-152286NB-I00 and CEX2020-001088-M) funded by MICIU/AEI/10.13039/501100011033 and by ERDF/EU; and the local government of Andalucia (reference PY20_00006). SM was supported by HORIZON-MSCA-2022-DN (grant no. 101120562, ProgRET) and JMR by the Juan de la Cierva program from the Spanish Ministry of Science, Innovation and Universities (AEI) (reference FJC2021-046858-I).

## Author Contribution

Conceptualization: JRMM, JMR

Methodology: JMR, SN, JT, RP

Investigation: JMR, RP, AGC, JC, SN, SM, ASO, AFM, JRMM

Visualization: JMR, JRMM

Supervision: JRMM, JMR

Writing—original draft: JMR, JRMM

Writing—review & editing: JRMM, JMR, JT

## Data and Materials availability

Datasets are available in the Gene Expression Omnibus (GEO) repository under the following accession numbers (GSE316738; GSE316739, and GSE318207).

Supplementary code has been deposited at GitHub as follows: Custom analysis scripts used in this study are available at https://github.com/rendon92/Macho_paper_2026. The MatchaiRen pipeline used for gene regulatory network inference is available at https://github.com/rendon92/MatchaiRen_v0.1. All other relevant data supporting the key findings of this study are available in the main text or the supplementary materials.

## Supplementary Figures

**Supplementary Figure S1.**
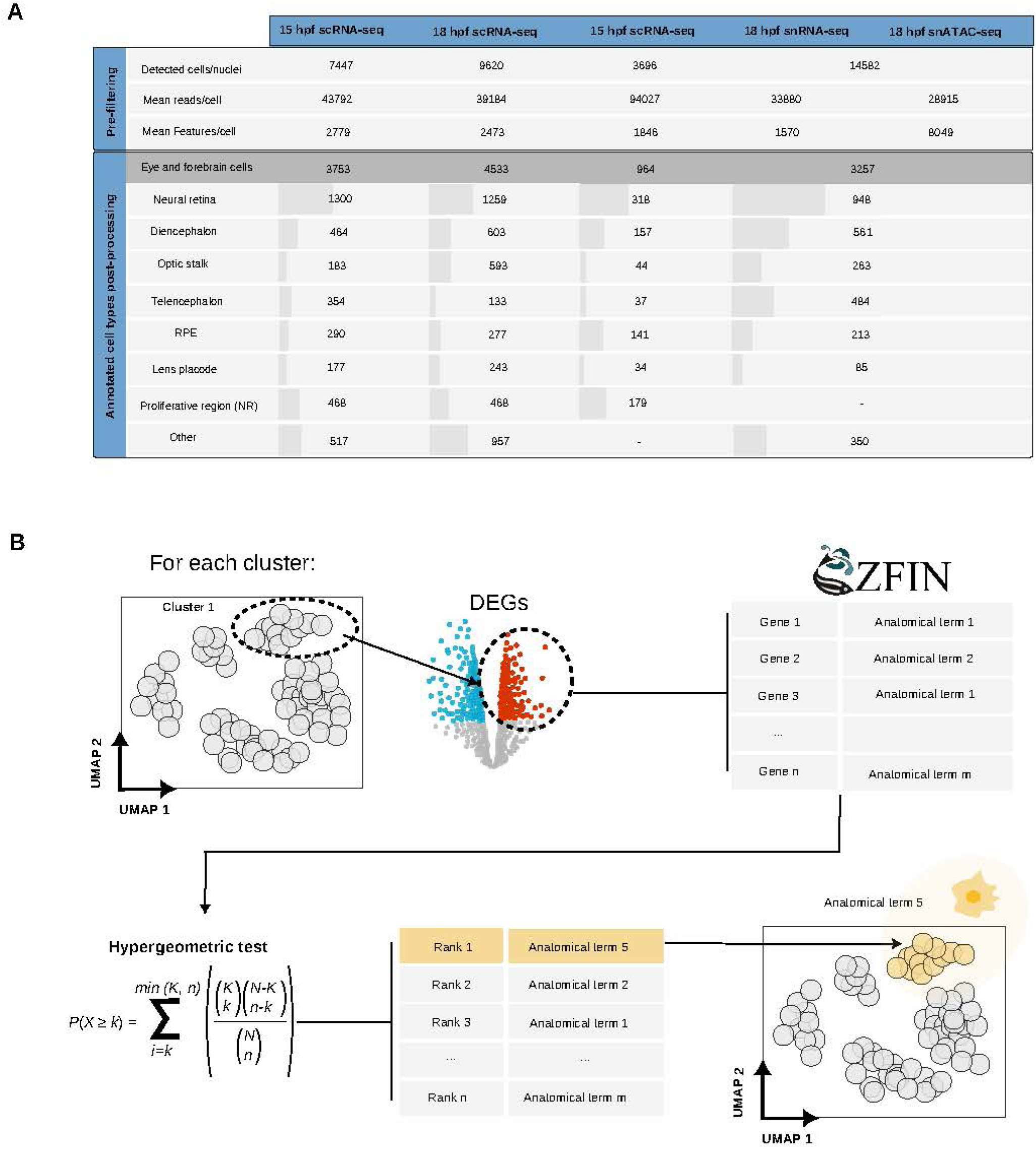
Dataset quality metrics and anatomical annotation pipeline. **(A)** Table summarizing QC metrics for scRNA-seq datasets at 15, 18 and 23 hpf and for the 18 hpf multiome experiment (snRNA-seq + snATAC-seq). Pre-filtering metrics include the total number of detected cells or nuclei, mean reads per cell/nucleus and mean number of detected features. After processing and annotation, number of cells annotated as “eye and forebrain” populations are reported, together with their distribution into neural retina, RPE, optic stalk, telencephalon, diencephalon, lens placode, the proliferative retinal compartment and additional domains. **(B)** Pipeline used to assign anatomical identity to clusters. For each dataset, cluster-specific marker genes were obtained with *FindAllMarkers*. These marker lists were intersected with ZFIN anatomical annotations filtered for genes expressed in the dataset. Enrichment for each anatomical term was quantified using a hypergeometric test. P-values were FDR-corrected and terms ranked by significance. The most enriched term was assigned as the cluster identity.

**Supplementary Figure S2.**
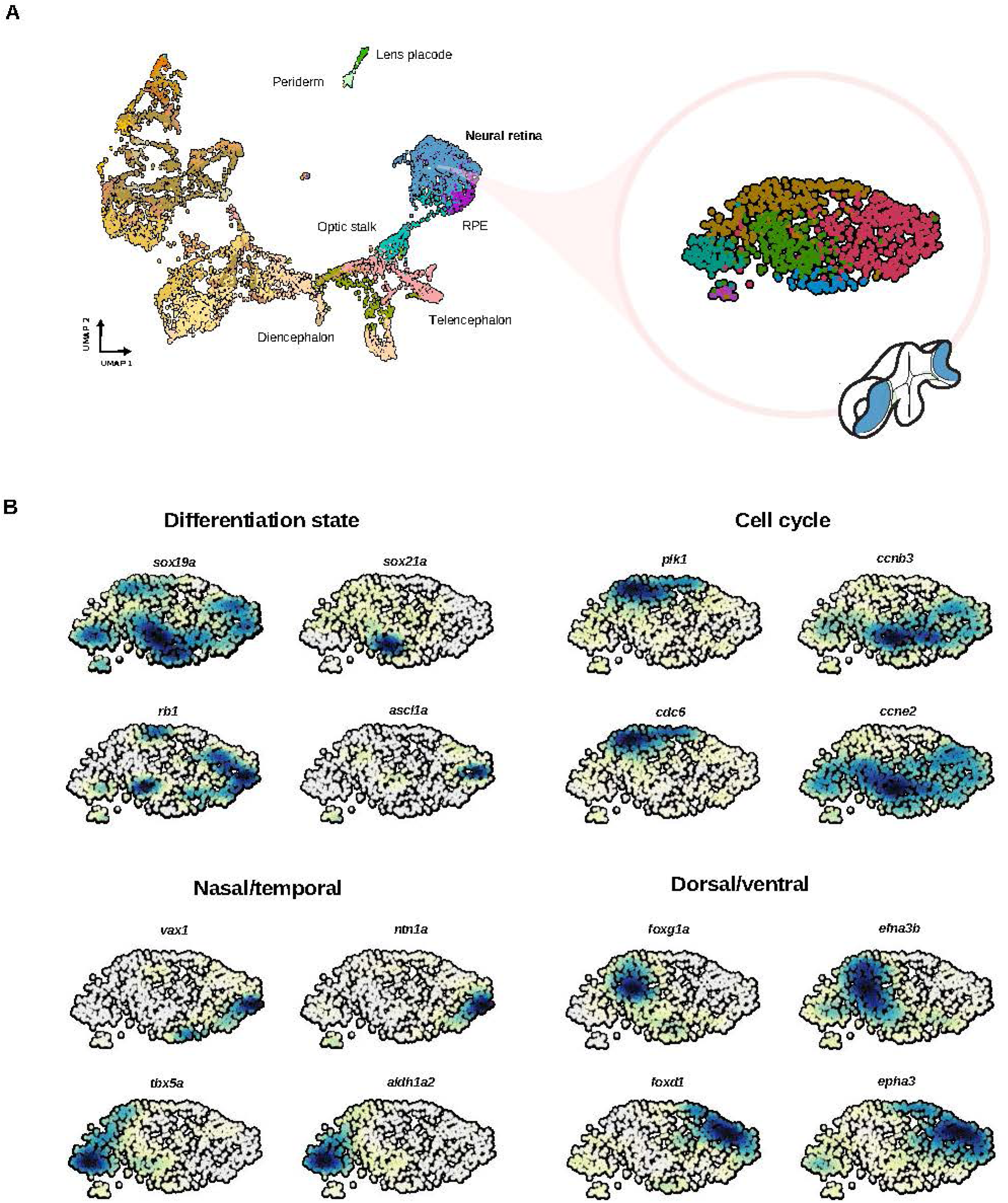
Retinal substructure and spatial markers in the 18 hpf multiome dataset. **(A)** Zoom-in of the 18 hpf multiome UMAP (from Fig. 3) highlighting the neural retina cluster. A schematic 3D reference, with the neural retina domain highlighted in blue, illustrates its spatial position within the eye primordium. **(B)** Density plots showing the expression of markers of differentiation state (*sox19a*, *sox21a*, *ascl1a*, *rb1*), cell cycle (*plk1*, *ccnb3*, *cdc6*, *ccne2*), and spatial axes (*tbx5a*/*aldh1a2* for dorsal; *vax1*/*ntn1a* for ventral; *foxg1a*/*efna3b* for nasal; and *foxd1*/*epha3* for temporal). Their distribution resembles the patterns observed in the single-cell integrated analysis (Figure 1D).

**Supplementary Figure S3.**
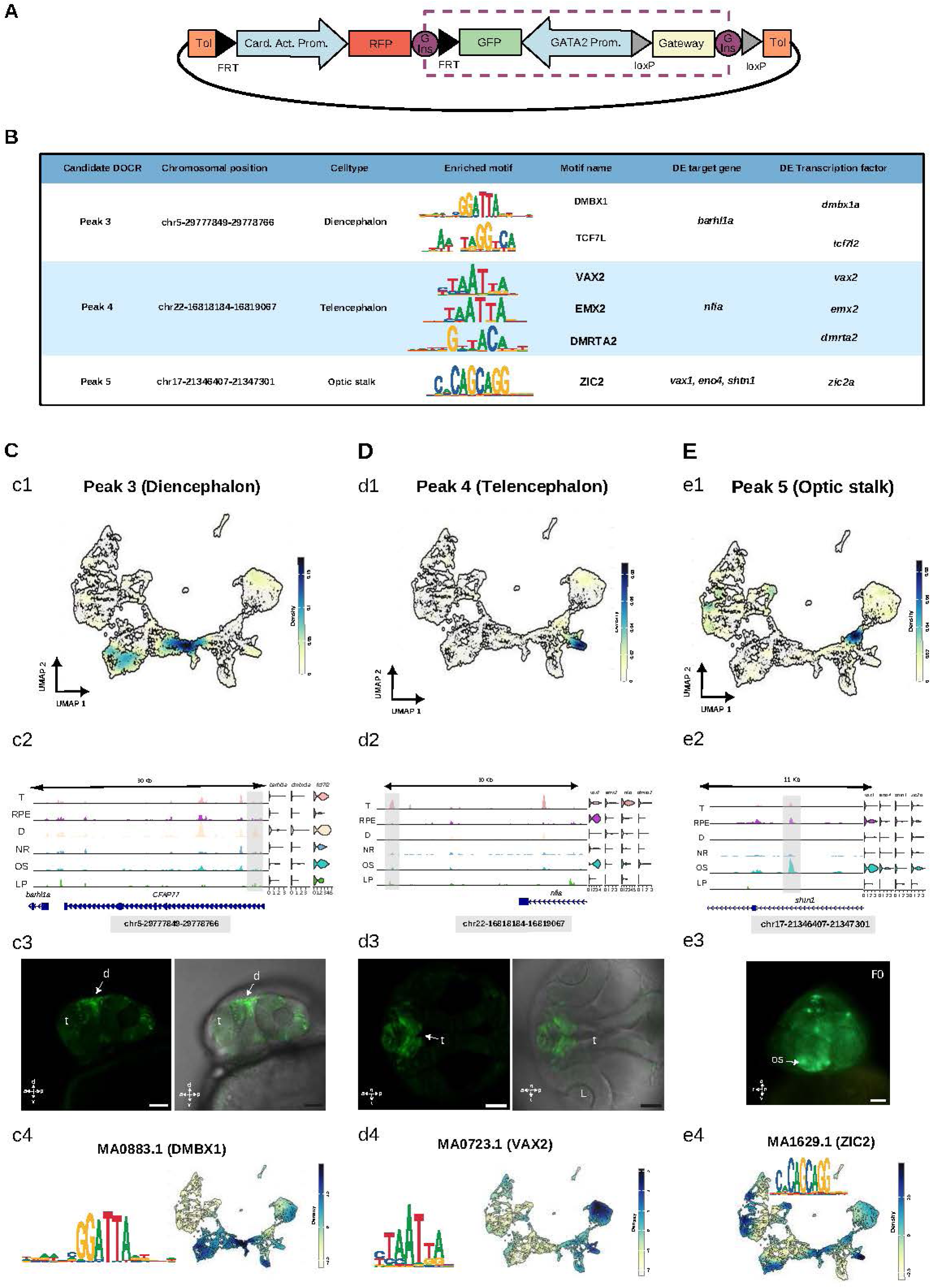
*In vivo* validation of additional domain-specific CREs. **(A)** Schematic representation of the ZED reporter vector used for cis-regulatory element assays (adapted from Bessa et al., 2009). **(B)** Summary table of three additional candidate CREs identified for dorsal diencephalon, telencephalon, and optic stalk. **(C)** Validation of the dorsal diencephalon CRE (chr5:29777849–29778766). **(c1)** Chromatin accessibility density plot showing strong enrichment within the diencephalic domain. **(c2)** Genomic view of the DOCR and nearby peaks; violin plots show expression of the proximal DEG *barhl1a,* as well as of the DEG *dmbx1a* and *tcf7l2*, encoding for TFs whose motifs are included in the region. **(c3)** Reporter expression in F1 embryos reveals specific GFP signal in dorsal diencephalon. **(c4)** Motif logo for DMBX1 (MA0883.1) and corresponding motif-activity distribution, including enriched activity in the diencephalon. **(D)** Validation of the telencephalic CRE (chr22:16818184–16819067). **(d1)** Chromatin accessibility density plot showing domain-specific signal in the telencephalic cell cluster. **(d2)** Genomic landscape surrounding the DOCR, with marked accessibility in telencephalon; violin plots show expression of *nfia,* as well as of the DEGs *vax2*, *emx2*, *dmrta2*, encoding for TFs whose motifs are present in the region. **(d3)** F1 reporter expression confirms activity restricted to the telencephalon. **(d4)** Motif logo for VAX2 (MA0723.1) and motif-activity UMAP highlighting strong activity in eye and anterior forebrain territories. **(E)** Validation of the optic stalk CRE (chr17:21346407–21347301). **(e1)** Chromatin accessibility density plot showing enrichment in the optic stalk. **(e2)** Genomic view of the region, with high accessibility in optic stalk; violin plots show expression of nearby DEGs (*vax2*, *eno4*, *shtn1*), a synexpression group for this domain, and of the DEG *zic2* whose motif is present in the region. **(e3)** F0 reporter embryos exhibit GFP expression consistent with optic stalk identity. **(e4)** Motif logo for ZIC2 (MA1629.1) and motif-activity distribution showing strong activity in the optic stalk region.

**Supplementary Figure S4.**
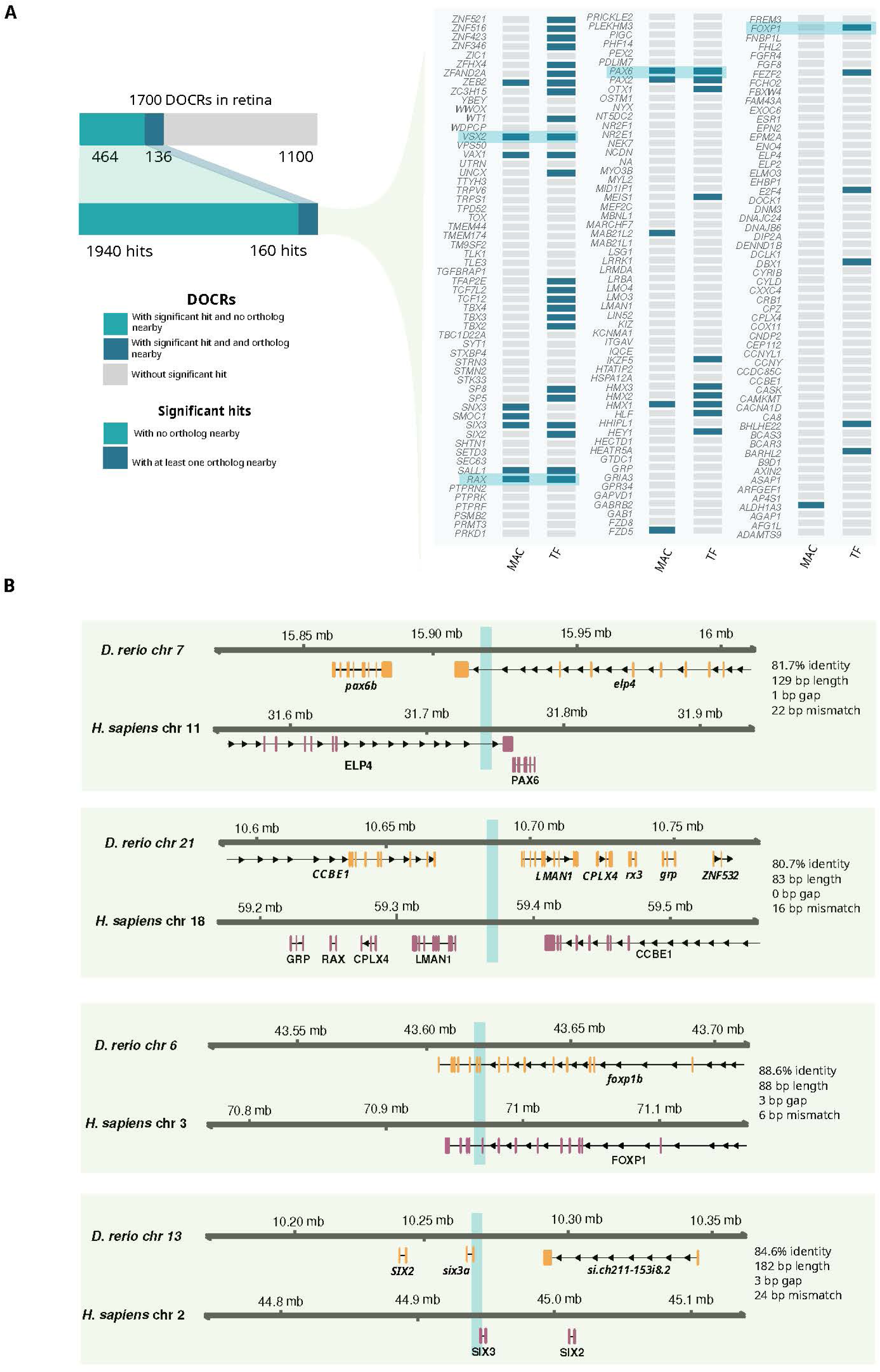
Conservation of zebrafish retinal DOCRs in the human genome. **(A)** Conservation analysis of 1,700 neural retina DOCRs (log₂FC ≥ 1, FDR ≤ 0.05). DOCRs are distributed into three categories: non-conserved (1,100 regions), conserved without proximity to orthologous gene pairs (464 regions; 1,940 total hits), and conserved near orthologous loci in both species (136 regions; 160 hits). The conserved sets (464 + 136) are magnified in a second chart. The 136 orthology-associated DOCRs correspond to 170 human genes, whose properties are summarized in a list indicating transcription factor identity and known association with MAC (microphthalmia/anophthalmia/coloboma) syndromes. **(B)** Representative examples of conserved CREs near shared orthologous genes in zebrafish and human. Conserved DOCRs at loci corresponding to relevant eye-field regulators and signaling components such as *PAX6/pax6b*, *RAX/rx3*, *FOXP1/foxp1b*, and *SIX2/SIX3* (zebrafish *six3a* and *six2*) are shown.

**Supplementary Figure S5.**
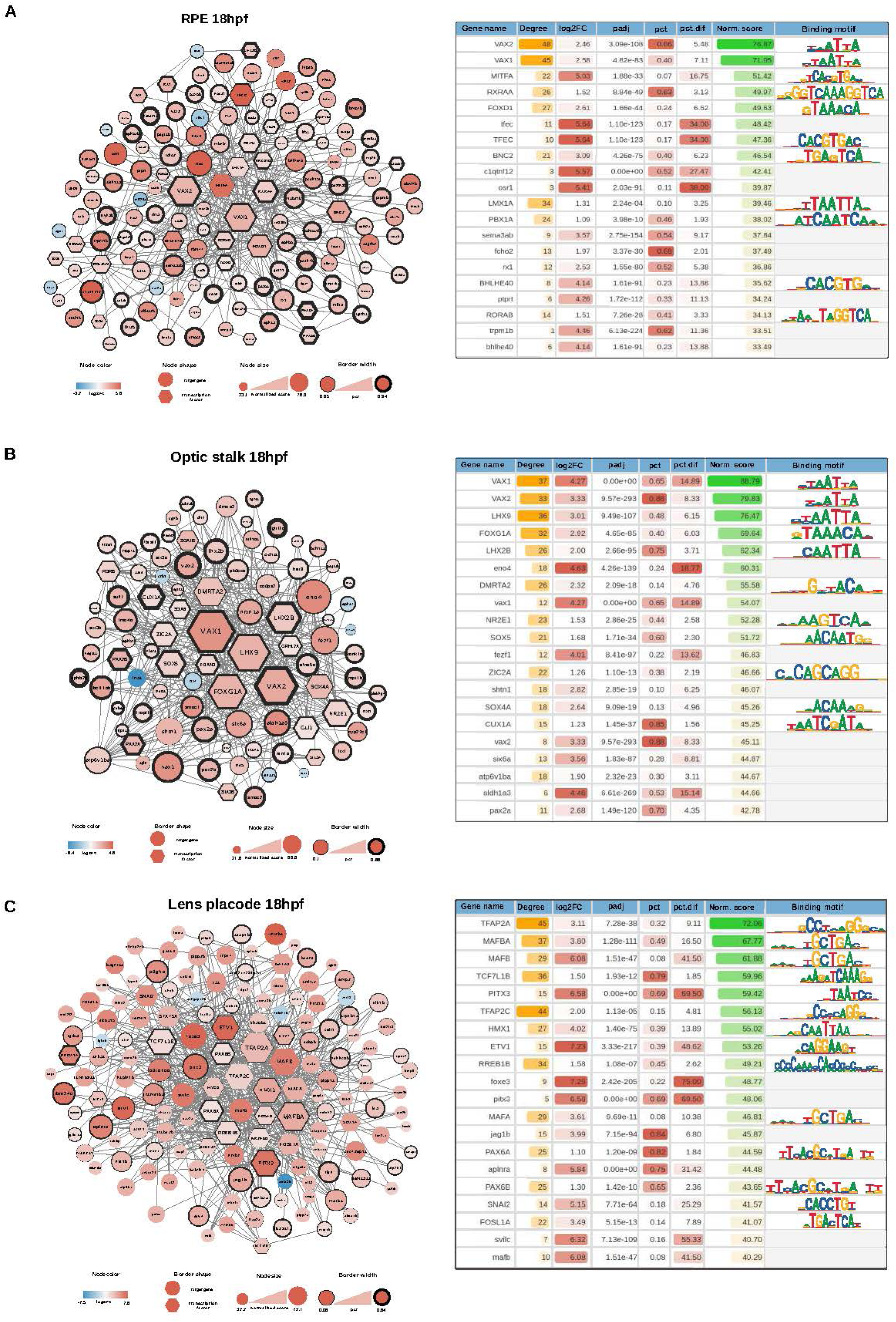
Gene regulatory networks inferred for RPE, optic stalk and lens placode domains at 18 hpf. **(A)** RPE network. The analysis highlights *vax1* and *vax2* as the most prominent regulators, together with genes encoding for BHLH-family TFs (*mitfa*, *tfec*, *bhlhe40*). The corresponding table lists the top 20 highest-scoring nodes based on connectivity and expression. **(B)** Optic stalk network. Top-ranked regulators include *vax1*, *vax2*, *lhx9* and *lhx2b*, alongside additional contributors such as *foxg1a*, *nr2e1*, *sox5*, and *zic2a*, outlining the main regulatory signals active in the optic stalk. **(C)** Lens placode network. TFAP and MAF families appear as key regulators, with *tfap2a*, *tfap2c*, *mafba*, *mafb*, *mafa*, and *pitx3* among the top ranked nodes.

**Supplementary Figure S6.**
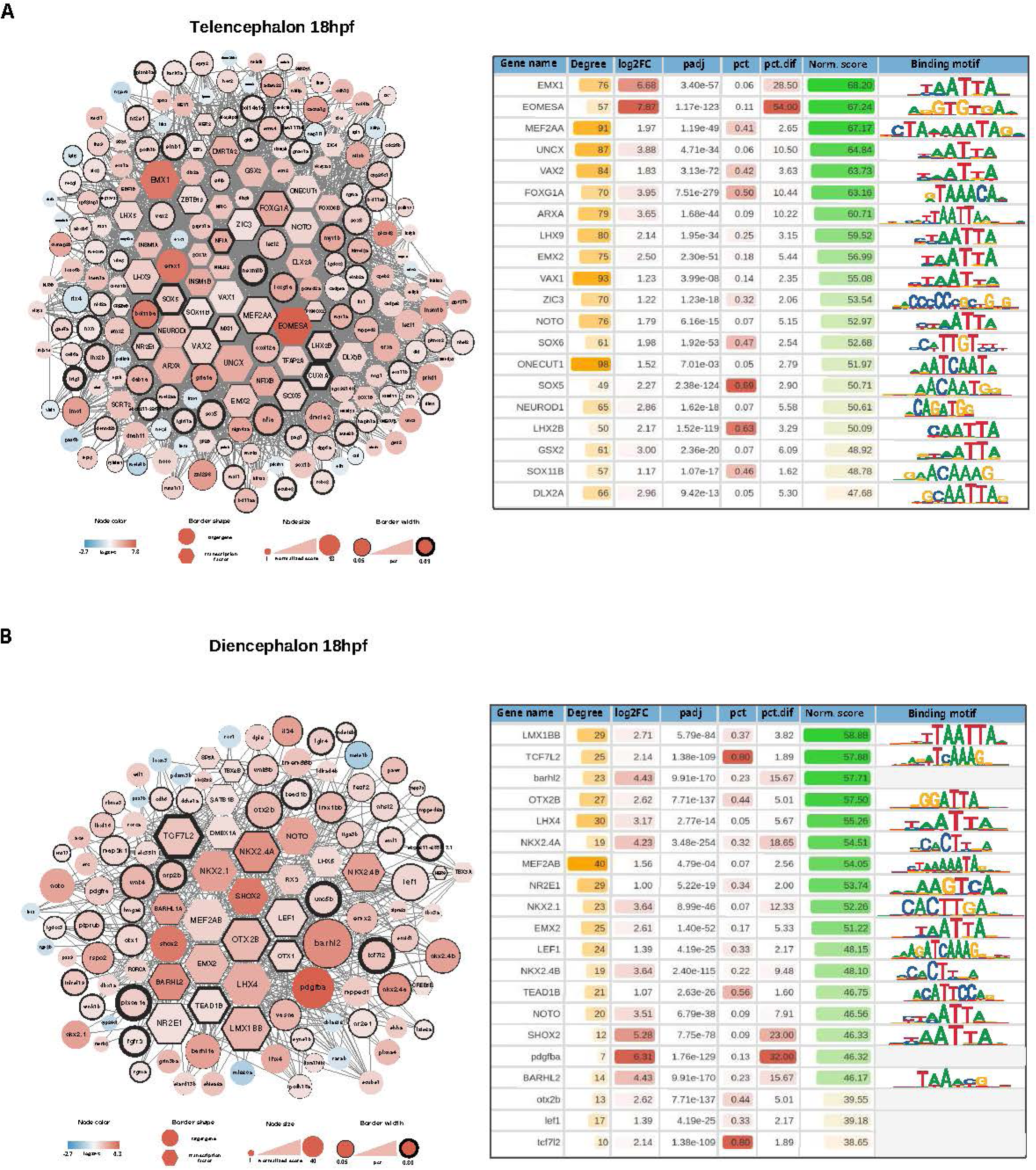
Gene regulatory networks inferred for telencephalon and diencephalon domains at 18 hpf. **(A)** Telencephalon network. Highest-scoring nodes include *emx1*, *emx2*, *eomesa*, *mef2aa*, *foxg1a* and several SOX-family genes, which together capture the main signals active in early telencephalic cells. **(B)** Diencephalon network. Key regulators include *lmx1bb*, *otx2b*, *nkx2.1*, *nkx2.4a/b*, together with *tcf7l2*, *barhl2*, *lef1* and *mef2ab*. These TF represent the major regulatory hubs active in the diencephalic domain at this stage.

**Supplementary Figure S7.**
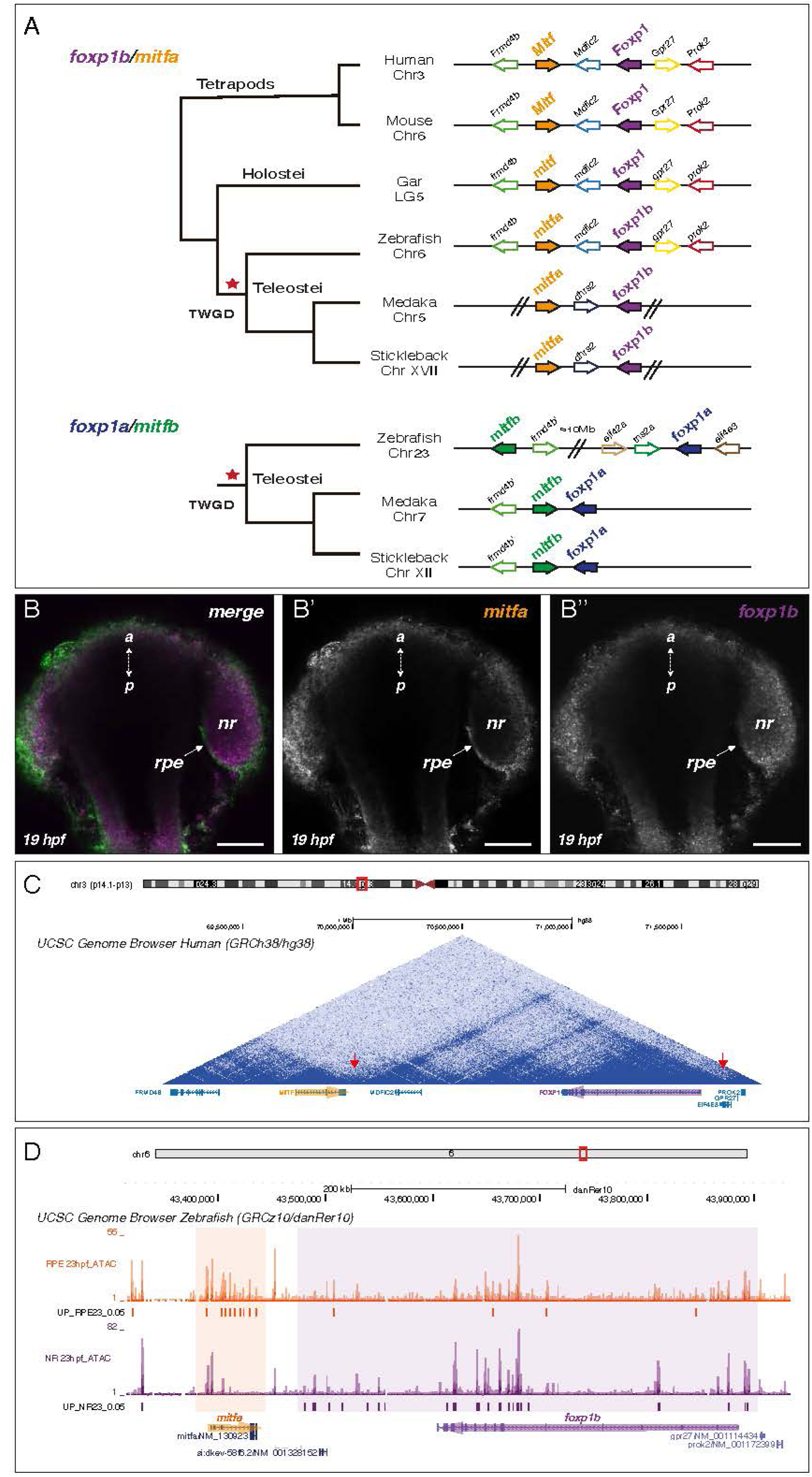
Analysis of conserved synteny between Foxp1 and Mitf. (A) The scheme depicts the syntenic arrangement of *Mitf* (orange/green) and *Foxp1* (purple/blue) ohnologs in the genomes of representative vertebrate species including tetrapods, holostei and teleosts. The corresponding chromosomes and duplicated paralogs in Teleostei are indicated. (B) HCR analysis of *mitfa* (B’) and *foxp1b* (B’’) expression patterns in 19 hpf zebrafish embryos shows complementary expression patterns at the RPE and neural retina (nr) domains respectively. a/p = antero-posterior; bar = 100 µm. (C) Micro-C track (UCSC genome browser) covering the *MITF*/*FOXP1* locus in the human genome (Chr 3) shows that *MITF* (orange) and *FOXP1* (purple) are included in neighboring TADs separated by a strong insulator boundary (red arrow). (D) 23hpf RPE and neural retina ATAC-seq tracks covering the *mitfa*/*foxp1b* locus (Chr 6) in the zebrafish genome (UCSC genome browser). Differentially opened chromatin regions (DOCRs) are indicated for the RPE (individual orange boxes) and neural retina samples (individual purple boxes). Note that the density of tissue specific DORCs define complementary cis-regulatory landscapes (shadowed large boxes) for *mitfa* and *foxp1b*.

**Supplementary Figure S8.**
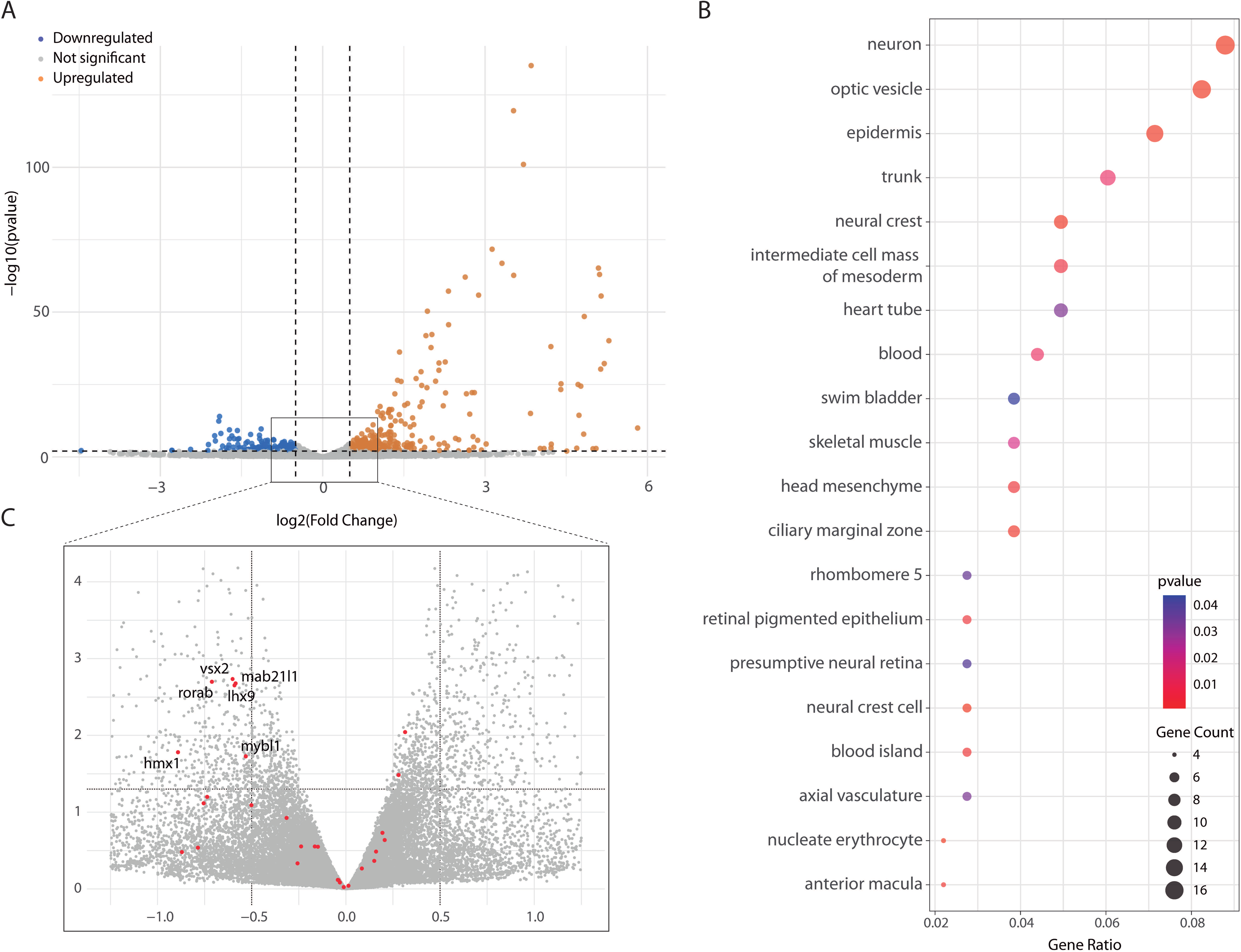
Transcriptomic analysis of zebrafish foxp1b crispants. (A) Volcano plot graph showing differentially expressed genes (DEGs) between wild type and *foxp1b* crispants at 18hpf. (B) Gene Ontology (GO) enrichment for downregulated genes in *foxp1b* crispants. (C) Overlay (red dots) of predicted *foxp1b* targets (i.e. MatchaiRen) on DEGs in the crispants. Note that several central nodes of the retinal network, predicted as *foxp1b* targets, are significantly downregulated in the crispants.

## List of supplementary Datasets description

**Supplementary Dataset 1. Differentially expressed genes across eye and forebrain domains in scRNA-seq data**

Differentially expressed genes (DEGs) identified in single-cell RNA-seq datasets of zebrafish embryos at 15, 18, and 23 hpf. Each row includes gene name, log2 fold-change, p-value, adjusted p-value, cellular prevalence, cluster identity, and sample origin. These data support domain annotation and subcluster analysis presented in Figure 1B–D.

**Supplementary Dataset 2. Top transcription factors expressed in each domain.**

List of the ten most significant transcription factors per domain based on p-value, along with their expression levels in each scRNA-seq dataset. Zebrafish orthology and previously reported domain expression annotations (ZFIN) are included. This dataset validates the domain-specific transcriptional programs described in the Results section.

**Supplementary Dataset 3. Differentially expressed genes (DEGs) and differentially open chromatin regions (DOCRs) in 18 hpf multiome data.**

Dataset containing DEGs and DOCRs identified for each annotated domain from single-nucleus multiome sequencing (snRNA-seq + snATAC-seq) of 18 hpf zebrafish embryo heads. Each domain is represented in a separate tab, listing transcriptionally and chromatin-accessibility significant features, including p-values, log2 fold-changes, and other standard metrics. These data link transcriptional programs to domain-specific regulatory landscapes, supporting the analyses shown in Figure 3A–C.

**Supplementary Dataset 4. Motif enrichment analysis for domain-specific regulatory programs in 18 hpf multiome data.**

Table of enriched motifs in domain-specific DOCRs at 18 hpf. For each major domain (neural retina, RPE, lens placode, optic stalk, telencephalon and diencephalon), the dataset lists motifs corresponding to TFs upregulated in the same domain, including observed counts, fold enrichment, p-values, and zebrafish orthologs (See Figure 3D–E).

**Supplementary Dataset 5. Conserved retinal DOCRs with proximal orthologs between zebrafish and human.**

Dataset of the sequence alignments (‘hits’) between zebrafish and human corresponding to zebrafish retinal DOCRs with proximal ortholog genes for both organisms. For each hit, the dataset lists zebrafish and human coordinates, associated genes and ortholog matches, highlighting conserved cis-regulatory regions linked to retinal regulators such as PAX6, RAX, SIX3, VAX1, VSX2, and OTX1 (See Supplementary Figure 4).

**Supplementary Dataset 6. Ranked gene regulatory network nodes for all domains inferred with MatchaiRen.**

This dataset contains the ranked node lists for the six domain-specific gene regulatory networks inferred with MatchaiRen (retina, RPE, optic stalk, lens placode, telencephalon, and diencephalon). For each domain, a table provides all network nodes together with their connectivity degree, differential expression statistics, cellular prevalence, and the normalized score used for ranking (see Methods section “Gene regulatory network inference with MatchaiRen” and Figure 5).

**Supplementary Dataset 7. Bulk RNA-seq analysis of *foxp1b* CRISPR mutants.**

This dataset compiles the results of the bulk RNA-seq experiment performed in *foxp1b* CRISPR-injected embryos to assess transcriptomic changes in the developing neural retina. It includes: a) A table summarizing the most significant differentially expressed genes (DEGs) obtained with DESeq2. b) A full list of all genes tested in the analysis. c) GO enrichment results for upregulated genes using anatomical ontology terms from ZFIN. d) GO enrichment results for downregulated genes using anatomical ontology terms from ZFIN. e) A list of predicted *foxp1b* retinal targets inferred with MatchaiRen, representing putative regulatory interactions.

**Supplementary Dataset 8. FACS gating strategy for isolation of eye-field and forebrain cells.**

Gating strategy used to isolate anterior neural plate–derived cells for scRNA-seq, including debris removal, singlet selection, live/dead discrimination using 7-AAD, and relaxed GFP-based gates applied to Tg(rx3:Gal4;UAS:GFP) embryos. This workflow corresponds to the procedure described in Methods “Embryo dissociation and single-cell libraries preparation”.

**Supplementary Dataset 9. CRISPR editing and CRE transgenesis.**

*Worksheet a*: Specific oligonucleotides used as a template in a *fill-in* PCR with the SpCas9 gRNA scaffold to synthetize the double stranded DNA containing the T7 promoter for subsequent *in-vitro* transcription of the single guide RNA sequence (sgRNA). Two different sgRNA pairs were designed to test whether they generated the same phenotype and to avoid native protein mRNA rescue experiments.

*Worksheet b*: Specific oligonucleotides used to detect DNA deletions flanking CRISPR target sites for Foxp1b in zebrafish and medaka. Wild-type amplicon size and optimum temperature for annealing step during PCR reactions are also included.

*Worksheet c*: Position of selected putative activating non-coding *cis*-regulatory elements (enhancers) in the zebrafish genome identified through a single-cell ATAC-seq in zebrafish embryos at 18hpf from retinal pigmented epithelium (RPE), neural retina (NR), optic stalk, lens, telencephalon, and diencephalon territories. Nearby upregulated genes located within 100kb upstream or downstream were identified via single-cell RNA-seq. Specific oligonucleotides, amplicon size and, optimum annealing temperature and cycles, for subsequent purification and cloning are also included.

